# Nanoscale 3D profiling of the T cell membrane reveals CD2 enrichment at microvilli tips, positioning adhesion near TCR zones in the immunological synapse

**DOI:** 10.64898/2026.08.17.744600

**Authors:** Suzi Kim, Elianna Lai, Aleksandra Klimas, Hamidreza Akhbariyoon, Emma F. DiBernardo, Yongxin Zhao, En Cai

**Affiliations:** Department of Biological Sciences, Carnegie Mellon University, Pittsburgh, PA, USA; Magnify Biosciences, Pittsburgh, PA, USA

**Keywords:** T cell, microvilli, immunological synapse, expansion microscopy, T cell receptor, adhesion receptor

## Abstract

The T cell membrane features a specialized molecular and topological organization critical for signaling and immune function. During antigen detection, T cells utilize microvilli to scan the antigen-presenting cell (APC) surface. Enriched with T cell receptors (TCRs) and key signaling molecules, microvilli serve as primary signaling hubs, yet their nanoscale architecture remains poorly defined. Upon antigen engagement, TCR activation drives the formation of the immunological synapse (IS). However, the IS nanoscale architecture remains technically challenging to profile. Here, **NanoMAP** (<u>Nano</u>scale <u>M</u>embrane <u>A</u>rchitecture <u>P</u>rofiling), an expansion microscopy-based platform, resolves receptor organization on T cell microvilli and within the IS at 35-60 nm resolution. Mapping the adhesion receptors CD2 and LFA-1 on OT-I T cells reveals that CD2 exhibits a stronger preference for microvillar tips than TCR and LFA-1. Within microvilli, CD2 displays a strong spatial association with TCR, and TCR and CD2 co-occupy close membrane contacts at the IS. After 1-2 h of antigen stimulation, TCR patches are selectively lost from microvilli, whereas CD2 and LFA-1 are retained. These results suggest a receptor-specific organization of adhesion receptors on microvilli and at the IS. Together, NanoMAP establishes a powerful framework for dissecting 3D nanoscale membrane architecture and spatial signaling in T cells.

**Key Results:**

- **NanoMAP** platform: Enables volumetric 3D nanoscale mapping and quantitative characterization workflow of membrane topology and receptor organization across whole T cells and immunological synapses (IS) between T cell-APC conjugates at 35-60 nm resolution.
- Distinct membrane profiles of OT-I CD8⁺ T cells at different states:

- Effector T cells possess a 5.1-fold higher microvillar density than naïve T cells.
- TCR patches are selectively lost from microvilli on effector T cells within 2 h of antigen stimulation.
- Differential receptor organization on effector OT-I T cell microvilli:

- CD2 patches are highly enriched in microvilli.
- Although TCR and LFA-1 patches are distributed across the majority of microvilli, they exhibit lower microvillar enrichment compared to CD2.
- Microvilli-associated CD2 patches show a strong spatial association with TCR patches.
- Immunological synapse (IS) nanoscale organization:

- Effector OT-I T cells and dendritic cells form a multifocal synapse.
- Within the IS, CD2 resides within TCR-proximal sites, whereas LFA-1 is excluded.

## Introduction

T cells are a critical component of the immune system. They survey host cells for signs of infection or malignancy and mount immune responses to eliminate them. Mechanistic insight into T cell function has driven the development of immunotherapies that have revolutionized cancer treatment, with dozens of FDA-approved treatments to date and thousands more in clinical trials.^[1–4]^ Despite these advances, the molecular mechanisms governing T cell activation and its regulation remain incompletely understood.^[5]^

At the cellular level, T cell activation and function require close membrane contact with antigen-presenting cells (APCs), enabling interactions between T cell surface receptors and their ligands that support antigen recognition, intracellular signaling, and execution of effector functions.^[6,7]^ Activation is triggered when the T cell receptor (TCR) binds its cognate peptide-major histocompatibility complex (pMHC) on the APC surface. TCR engagement initiates a downstream signaling cascade that drives formation of the immunological synapse (IS), a tight cell-cell contact with the APC.^[8]^ Within the synapse, TCRs rapidly form microclusters that move to the center to form the central supramolecular activation cluster (cSMAC), while the adhesion receptor LFA-1 binds to ICAM-1 on the APC and translocates to the periphery to form the peripheral supramolecular activation cluster (pSMAC). This highly organized structure is essential for sustaining T cell activation and enabling efficient function.^[7,9,10]^ As such, the T cell membrane and the modulation of its nanoscale organization serve as the central platform for T cell immune responses, highlighting the importance of understanding their molecular basis and regulation.

T cells are highly sensitive and can respond to as few as several molecules of pMHC, and recent evidence suggests that the nanoscale organization of their membrane architecture, including receptor distribution and surface topology, facilitates antigen detection and signal initiation.^[11–15]^ Recent data show that T cells pre-organize surface receptors, including TCRs, and signaling molecules into nanoscale clusters prior to activation. This increases TCR signaling strength and enables rapid assembly of TCR microcluster signalosomes.^[16–19]^

In addition, the T cell surface features a dense coat of finger-like membrane protrusions called microvilli, which are highly dynamic and survey the APC surface within one minute.^[20,21]^ Microvilli tips establish tight contacts with the antigen-presenting surface and bring TCRs within 15 nm of pMHC, promoting TCR-pMHC interactions.^[15,22,23]^ Furthermore, TCRs and several key signaling molecules, including CD2, CD4, and Lck, are found to be enriched within microvilli, further promoting TCR signaling at these sites. These results support microvilli as signaling hubs for efficient antigen detection and initiation.^[24–26]^ However, exactly how these molecules are organized within microvilli remains unclear. Notably, T cells engage APCs through distinct adhesion receptors at different levels of membrane contact, yet it remains unclear how these receptors are organized on microvilli, which serve as initial contact points, and how this organization is remodeled during synapse formation. More broadly, capturing the 3D membrane architecture and nanoscale receptor organization across the entire T cell surface remains a major technical challenge — particularly within the IS, where the synaptic cleft is only tens of nanometers wide.

In this work, we employed expansion microscopy (ExM) to visualize the nanoscale architecture of the entire T cell membrane. ExM enables nanoscale imaging of biological samples by isotropically expanding the specimen after chemical processing, thereby achieving nanoscale resolution using conventional confocal microscopy.^[27–29]^ Here, we use ExM to map the entire 3D surface of T cells and examine the spatial correlations between microvilli and receptors at 60 nm resolution, and to visualize surface topology and receptor distribution within the IS at 35 nm resolution. In parallel, we developed a data analysis framework that quantifies the spatial relationships of receptor patches on microvilli across the cell, providing a 3D nanoscale map of the T cell’s membrane topology and receptor distribution. We further applied this approach to a panel of key surface receptors on T cells, mapping the nanoscale architecture of the T cell membrane. We term this platform T cell <u>Nano</u>scale <u>M</u>embrane <u>A</u>rchitecture <u>P</u>rofiling (**NanoMAP**).

With NanoMAP, we profiled the spatial organization of CD2, LFA-1, TCR and microvilli on effector CD8^+^ T cells from OT-I transgenic mice. CD2 and LFA-1 are the two major adhesion receptors on T cells that have distinct molecular properties and play different roles in T cell functions: CD2 is a costimulatory receptor that amplifies TCR signaling, whereas LFA-1 is an integrin that mediates firm adhesion.^[30,31]^ Our results show that CD2 is more enriched in microvilli relative to TCR and LFA-1, which are distributed in proportion to membrane. Within the IS, CD2 coexist with TCR in tight membrane contact areas where LFA-1 is excluded.

Together, these results highlight the differential nanoscale organization of TCR, CD2 and LFA-1 across the membrane of effector OT-I T cells at different activation stages, suggesting a receptor-specific nanoscale organization of the T cell membrane.

## Results

### Expansion microscopy workflow enables nanoscale profiling of the T Cell membrane architecture

To visualize nanoscale receptor patches and T cell microvillar architecture, we adapted the established Magnify ExM method reported by Klimas et al. to T cells and T cell-APC conjugates.^[29]^ All measurements reported in this work were conducted using effector OT-I T cells, referred to below as T cells or OT-I T cells, unless stated otherwise. Primary mouse T cells were first labeled with primary antibodies against receptors of interest, fixed, and then stained with fluorescent secondary antibodies. Briefly, fixed cells were suspended in a gelling solution and polymerized under UV light to generate a hydrogel in which cells and their proteins were covalently anchored. The gel was then treated with a homogenization solution that disrupts intermolecular crosslinks and mechanical constraints, enabling isotropic expansion while preserving cellular nanostructures. After homogenization, the gel was immersed in 1x PBS and allowed to expand to 4.3-fold of its original size.

Because the homogenization step involves harsh processing conditions, a substantial fraction of the antibody signal from pre-expansion labeling can be lost. The heat-induced denaturation of the antibody epitopes during homogenization and the dilution of fluorescence signal following the volumetric expansion are known contributors to reduced labeling efficiency in samples with only pre-expansion antibody labeling. To recover and enhance labeling, we performed a second round of antibody staining after expansion, as previously described by Klimas et al. and Zwettler et al.^[29,32]^ Post-expansion “rescue” staining has been shown to restore fluorescence intensity and access epitopes that were sterically inaccessible in the unexpanded state, thereby markedly improving labeling quality, especially for sparse or low-abundance molecules (**Figure 1a**). In this post-expansion staining, we also included fluorescently labeled wheat germ agglutinin (WGA) to label the plasma membrane.

**Figure 1.**
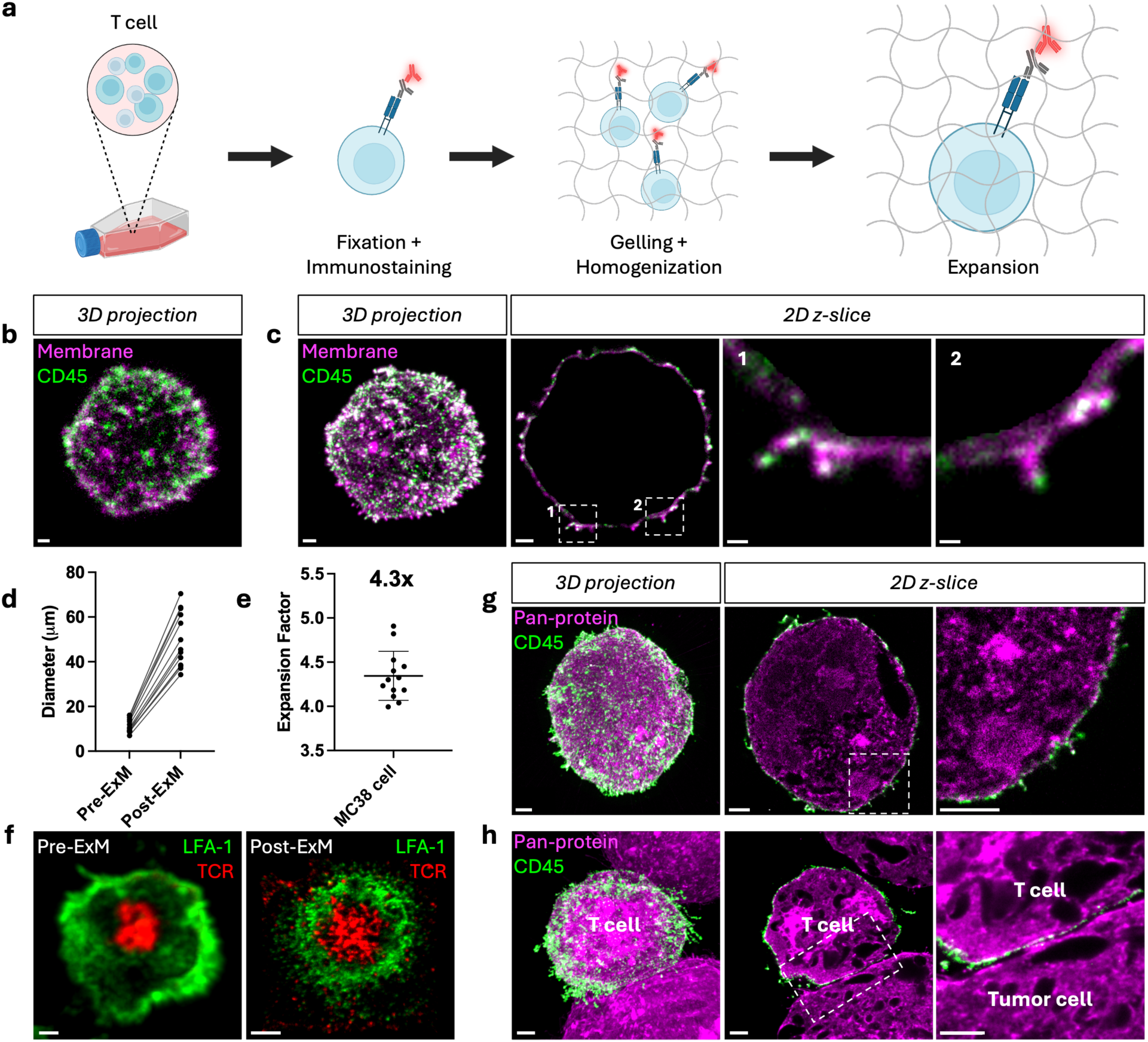
NanoMAP enables nanoscale visualization of whole T cell membrane with 60 nm resolution for T cell profiling. a) Schematic ExM workflow with T cells. b) 3D maximum projection image of a fixed OT-I T cell labeled with anti-CD45-AF488 (green) and WGA-CF633 (magenta). c) Post-expansion 3D maximum projection image and 2D z-slice image of the T cell from (b). d) Major axis diameter of MC38 tumor cell nuclei in pre-expansion and post-expansion in 1x PBS. e) Expansion factor in 1x PBS calculated by measuring the major axis diameter of MC38 tumor cell nuclei in pre-expansion and post-expansion. Sample sizes in (d-e): *n* = 13 cells from N = 3 gels. Data are presented as mean ± S.D. f) Pre-expansion and post-expansion images of T cell labeled with anti-TCR-AF594 (red) and anti-CD11a-AF488 (green) after 7.5 min contact with supported lipid bilayer. g) Representative image of OT-I T cell fully expanded in H_2O_ labeled with anti-CD45-AF488 and NHS-ester-Cy5 shown in 3D projection and 2D z-slice with a magnified image of the region within the white box. h) Representative image of OT-I T cell conjugated with MC38 tumor cell and expanded in H_2O_. T cell-MC38 conjugates are labeled with anti-CD45-AF488 (green) and NHS-ester-Cy5 (magenta) and shown in 3D projection and 2D z-slice with a magnified image of the region within the white box. Scale bar = 1 μm. Scale bar in magnified 2D images in (c) = 200 nm.

To benchmark the gain in effective resolution, we stained OT-I T cells for the abundant surface receptor CD45 together with membrane labeling and imaged the same cell before and after expansion under identical confocal settings. The pre-expansion image poorly resolved membrane topology and receptor patches (**Figure 1b**), whereas the post-expansion image clearly revealed sub-diffraction membrane protrusions and discrete CD45 patches (**Figure 1c**). To determine the expansion factor and convert post-expansion measurements to biological (pre-expansion) dimensions, we measured the major axis diameter of the same nuclei before and after expansion, yielding an expansion factor of 4.3-fold in MC38 tumor cells (**Figure 1d-e, Figure S1a-b**). For OT-I T cells, the expansion factor was calculated as the ratio of the mean post-expansion to mean pre-expansion nuclear diameter. To confirm that the expansion factor does not depend on T cell differentiation state, we measured the expansion factor of naïve and effector OT-I T cells separately. Both naïve and effector OT-I cells expanded 4.4-fold (naïve: 95% CI 4.3-4.6; effector: 95% CI 4.2-4.7), consistent with the 4.3-fold measured in MC38 tumor cells (**Figure S1c-d**).

Next, we assessed the level of local distortion introduced by expansion using the 2D distortion analysis described by Klimas et al.^[29]^ The root-mean-square (RMS) length-measurement error was calculated as a function of measurement length for two separate channels. The RMS error remained below 3% for measurement lengths up to 50 µm in both channels, comparable to the 2.5% reported by Klimas et al., indicating near-isotropic expansion with minimal local distortion (**Figure S1e-f**).

We next applied this workflow to visualize the IS, using OT-I T cells activated on supported lipid bilayers presenting the agonist pMHC (H-2K^b^–OVA^257-264^) and ICAM-1 to resolve nanoscale receptor organization at the interface. As expected, pre-expansion synapses displayed the classical “bull’s-eye” pattern, with TCR-enriched cSMAC surrounded by an LFA-1-rich peripheral ring (pSMAC). After expansion, the IS exhibited sharply delineated cSMAC and pSMAC boundaries and resolved individual receptor patches within cSMAC and pSMAC (**Figure 1f**).

To further increase effective resolution, we immersed the gel-embedded cells in water to achieve an approximately 7.3-fold linear expansion, 7.4-fold in Jurkat cells (95% CI 6.9-8.0) and 7.2-fold in OT-I T cells (95% CI 6.9-7.6), corresponding to ∼35 nm lateral resolution (**Figure S1g-h**). In this condition, T cells were labeled with anti-CD45 and a fluorescent NHS ester to mark total protein content. The resulting images (**Figure 1g**) revealed substantially finer cellular detail than those at 4.3-fold expansion (**Figure 1c**). Using this higher expansion, we visualized the membrane organization at the synaptic cleft between an OT-I T cell and an antigen-bearing tumor cell (MC38-OVA) (**Movie S1**). **Figure 1h** shows tight and loose contact regions within the synapse with CD45 exclusion from tight membrane contacts.

With ExM’s capability to examine the membrane structure at nanoscale, we compared the surface topology of naïve and effector T cells (**Figure 2a-b**, **Movie S2**). Naïve T cells displayed relatively smooth surfaces with few microvilli, whereas effector T cells exhibited a ∼5.1-fold higher microvillar density and a ∼2.8-fold larger cell surface area, indicating substantial remodeling of the plasma membrane as T cells transition between functional states (**Figure 2c-d**).

**Figure 2.**
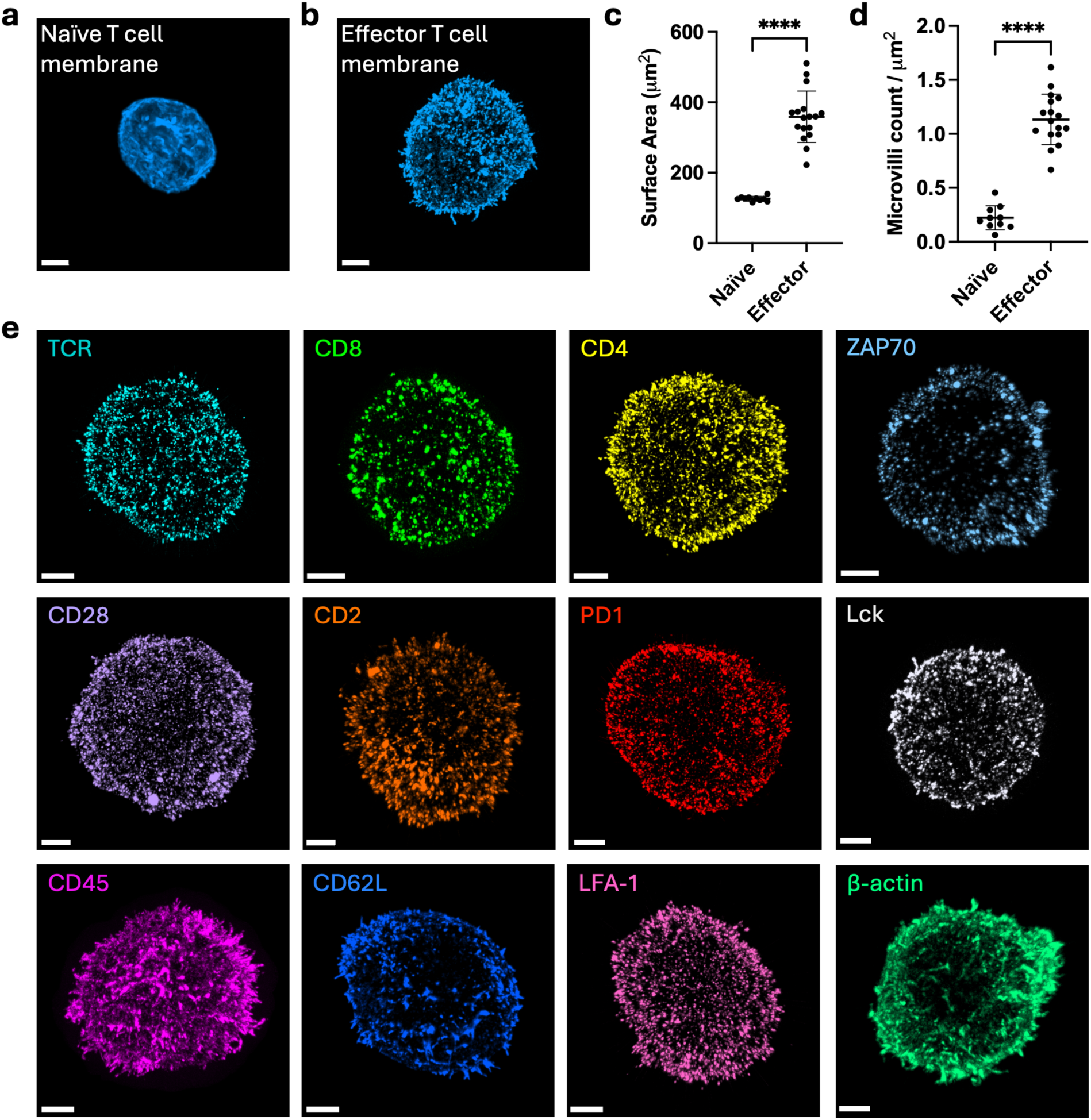
Profiling different T cell states and surface receptor organization using ExM. a-b) Post-expansion 3D projection image of (a) naïve OT-I T cell and (b) effector OT-I T cell labeled with WGA-CF633 (blue). c) Mean surface area per cell of naïve T cells and effector T cells (\*\*\*\**p* < 0.0001). d) Mean microvilli density of naïve T cells and effector T cells (\*\*\*\**p* < 0.0001). Data are presented as mean ± S.D. Statistical significance tested using unpaired two-tailed Welch’s t-test. Sample sizes: Naïve T cell (*n* = 10 cells from N = 1 mice); effector T cell (*n* = 17 cells from N = 4 mice). e) Post-expansion 3D projection image panel. From left to right: (top) TCR, CD8, CD4, ZAP70, (middle) CD28, CD2, PD1, Lck, (bottom) CD45, CD62L, LFA-1, β-actin. All images show effector OT-I T cells except CD4, which shows a CD4⁺ T cell isolated from C57BL/6 mice. Scale bar = 2 μm.

To further decode T cell membrane architecture, we sought to map the nanoscale distribution of key receptors that regulate activation. Although antibodies for various surface receptors are readily available, many are not compatible with ExM because the workflow can alter proteins into a non-native or partially denatured state and change their steric environment through hydrogel embedding. This can result in disrupted epitope accessibility and abrogate antibody binding. Consequently, a major bottleneck is identifying antibody clones that retain robust binding and labeling efficiency after the expansion workflow. Through systematic screening of commercially available antibodies, we established a panel targeting stimulatory, inhibitory, and adhesion receptors, as well as intracellular signaling proteins and the cytoskeletal protein β-actin, that are compatible with our ExM workflow, enabling nanoscale mapping of their spatial organization in T cells (**Figure 2e**). While subsequent experiments presented in this work were conducted using primary mouse T cells, we also demonstrated that this platform is compatible with primary human T cells (**Figure S1i**).

### Quantitative characterization of microvilli (MV) and receptor patches

To characterize nanoscale membrane topology and receptor organization, we developed a quantitative 3D analysis workflow that detects receptor patches and microvillar structures on the T cell surface and extracts their geometric and spatial properties. This in-house workflow operates on fully 3D volumetric data and is implemented using Imaris (v10.0.1, Bitplane). First, we segmented the cell using the membrane channel by adjusting intensity thresholds to include only the main cell body while excluding microvilli. Because our goal was to analyze membrane-localized receptors, we restricted the analysis to the defined cell surface regions and excluded intracellular fluorescence signals (**Figure S2a-b**). This procedure generated a 3D surface representing the boundary of the T cell body, which we used as a reference surface to define the cellular outline (**Figure S2c**). We then segmented membrane protrusions extending beyond this boundary as microvilli (**Figure 3a-b**). The volume associated with these protrusions was defined as the microvillus (MV) region (**Figure 3c**, white), while the remaining membrane was designated as the cell body region (**Figure 3c**, magenta). To validate the accuracy of this microvilli segmentation pipeline, we manually measured microvillus length and diameter and found close agreement with the automated measurements (**Figure S3**). We further confirmed that the segmented protrusions are microvilli by co-labeling OT-I T cells with WGA and an antibody against β-actin, which is the main component of the actin bundles that form the microvilli (**Figure S2d**). ^[33]^ Approximately 94% of the segmented microvilli contained segmented β-actin and β-actin occupied approximately 60% of the volume of WGA-defined microvilli (**Figure S2e-f**). These results confirm that our segmentation predominantly captures actin-based microvilli.

**Figure 3.**
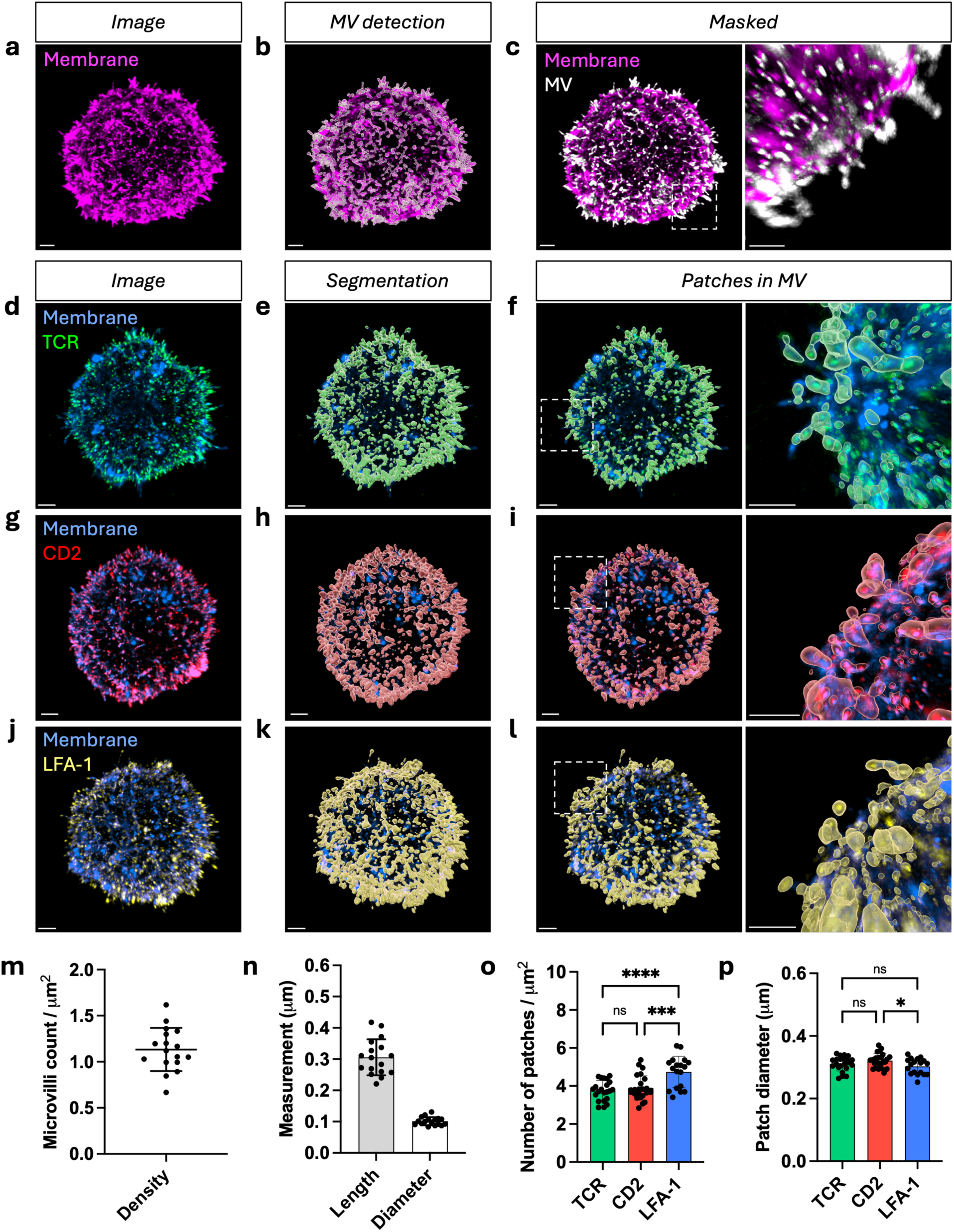
Microvilli (MV) and patch segmentation and quantification of 3D post-expansion images. a) Post-expansion 3D image of OT-I T cell labeled with WGA-CF633 for membrane staining. b) Image from (a) showing MV segmentation using Imaris. c) Image from (a) with the MV region in white and the cell body (non-MV region) in magenta. Magnified image of the region within the white box. d) Post-expansion 3D image of OT-I T cell labeled with anti-TCR-AF488 (green) and WGA-CF633 (blue). e) Image from (d) showing TCR patch segmentation. f) Image from (d) showing TCR patches localized in the MV region. Magnified image of the region within the white box. g) Post-expansion 3D image of OT-I T cell labeled with anti-CD2-AF555 (red) and WGA-CF633 (blue). h) Image from g) showing CD2 patch segmentation. i) Image from (g) showing CD2 patches localized in the MV region. Magnified image of the region within the white box. j) Post-expansion 3D image of OT-I T cell labeled with anti-CD11a-AF488 (yellow) and WGA-CF633 (blue). k) Image from (j) showing LFA-1 patch segmentation. l) Image from (j) showing LFA-1 patches localized in the MV region. Magnified image of the region within the white box. m) Mean microvilli density per cell calculated by number of microvilli per area. n) Mean microvilli length and diameter per cell. Sample sizes in (m-n): *n* = 17 cells from N = 4 mice. o) Mean patch density of TCR, CD2, and LFA-1 calculated by number of patches per area (CD2 vs. LFA-1: \*\*\**p* = 0.0004; CD2 vs. TCR: *p* = 0.7661; TCR vs. LFA-1: \*\*\*\**p* < 0.0001; ns = not significant). p) Mean diameter of TCR, CD2, and LFA-1 patches (CD2 vs. LFA-1: \**p* = 0.0350; CD2 vs. TCR: *p* = 0.2878; TCR vs. LFA-1: *p* = 0.5252; ns = not significant). Data are presented as mean ± S.D. In (o-p), statistical significance was determined using ordinary one-way ANOVA followed by Tukey’s multiple comparisons test. Sample sizes: TCR (*n* = 22 cells from N = 6 mice), CD2 (*n* = 24 cells from N = 6 mice), and LFA-1 (*n* = 18 cells from N = 4 mice). Scale bar = 1 μm.

By construction, the MV region identified in this way excludes the main cell body and captures only the protruding portion of each microvillus from its base. Very short, bud-like protrusions that do not extend beyond the cell-body membrane signal may not reach the detection threshold (0.15 μm; see Methods) from the reference surface (**Figure S2c**) and would therefore go undetected, leading to a slight underestimation of microvillus density. In addition, the segmentation procedure may exclude the proximal portion of protrusions that overlaps with the cell-body membrane signal. Thus, our analysis explicitly emphasizes more extended protrusions that are most likely to engage antigen-presenting cells (APCs) during the earliest stages of contact.

We next segmented receptor patches for TCR, CD2, and LFA-1 from their respective fluorescent channels (**Figure 3d-e, 3g-h, 3j-k**, **Figure S4a-c**). Here, we define receptor patches as regions of receptor signal above threshold for detection, which may contain one or more receptor molecules. Using the MV regions defined, as shown in **Figure 3c**, we then classified receptor patches according to whether they resided within microvilli or on cell body, allowing us to identify microvillus-associated patches across the entire T cell membrane (**Figure 3f, 3i, 3l**). It is important to note that patch detection depends on the local signal to noise ratio of fluorescence intensity; as a result, individual receptors and very small or dim patches may not be detected and are excluded from the analysis.

This integrated analysis workflow enables quantitative 3D mapping of both microvillar architecture and receptor organization on the T cell membrane **(Movie S3**). We quantified microvilli density, length, and diameter, as well as the density and size of TCR, CD2, and LFA-1 patches, which were consistent with previously reported measurements (**Figure 3m-p**).^[25,33,34]^

Together with our ExM-based super-resolution imaging workflow, this established a platform for profiling the nanoscale spatial distribution of specific receptors over the 3D membrane topology of T cells, which we term T cell **Nano**scale **M**embrane **A**rchitecture **P**rofiling (**NanoMAP**).

### CD2 is more enriched on microvilli relative to TCR and LFA-1

T cells express a diverse set of adhesion receptors that cooperate to promote membrane engagement with APCs and to tune TCR signaling.^[35]^ Here, we sought to define the nanoscale distribution of adhesion receptors on microvilli and their spatial relationship to TCR. We focused on two key adhesion receptors, CD2 and LFA-1, which have distinct molecular properties and nonredundant roles in T cell function (**Figure 4a-d**).

**Figure 4.**
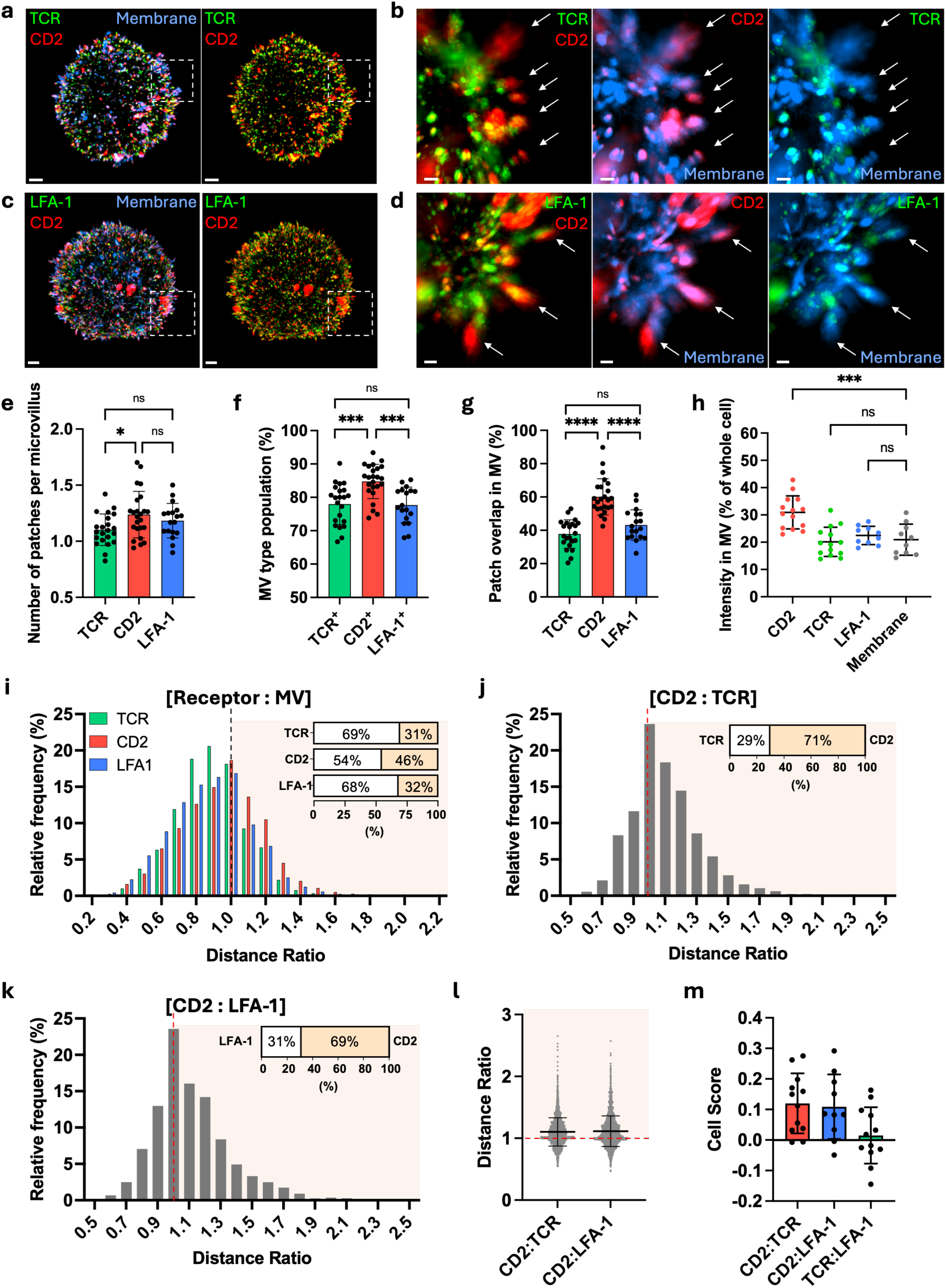
CD2 patches are enriched in MV region relative to TCR and LFA-1. a) Post-expansion 3D image of OT-I T cell labeled with anti-TCR-AF488 (green), anti-CD2-AF555 (red) and WGA-CF633 (blue) for membrane staining. b) Magnified image of the region within the white box from (a). c) Post-expansion 3D image of T cell labeled with anti-CD11a-AF488 (green), anti-CD2-AF555 (red) and WGA-CF633 (blue). d) Magnified image of the region within the white box from (c). In (b) and (d), white arrows are pointing at microvilli tips. e) Mean number of TCR, CD2, and LFA-1 patches localized per microvillus per cell (CD2 vs. LFA-1: *p* = 0.5757; CD2 vs. TCR: \**p* = 0.0259; TCR vs. LFA-1: *p* = 0.3018; ns = not significant). f) Population of microvilli containing TCR, CD2, and LFA-1 patches per cell (CD2 vs. LFA-1: \*\*\**p* = 0.0004; CD2 vs. TCR: \*\*\**p* = 0.0003; TCR vs. LFA-1: p = 0.9842; ns = not significant). g) Mean occupancy (voxel overlap) of TCR, CD2, and LFA-1 in the MV region per cell (CD2 vs. LFA-1: \*\*\*\**p* < 0.0001; CD2 vs. TCR: \*\*\*\**p* < 0.0001; TCR vs. LFA-1: *p* = 0.1865; ns = not significant). In (e-g), data are presented as mean ± S.D. and statistical significance was determined using ordinary one-way ANOVA with Tukey’s comparisons test. Sample sizes: TCR (*n* = 22 cells from N = 6 mice), CD2 (*n* = 24 cells from N = 6 mice), and LFA-1 (*n* = 18 cells from N = 4 mice). h) Percentage of intensity sum of CD2, TCR, LFA-1 and membrane staining in the MV region relative to the intensity sum in the whole cell. Data are presented as mean ± S.D. Statistical significance was determined using ordinary one-way ANOVA with Dunnett’s comparisons test (membrane vs. TCR: *p* = 0.9653; membrane vs. LFA-1: *p* = 0.8456; membrane vs. CD2: \*\*\**p* = 0.0001; ns = not significant). Sample sizes: TCR (*n* = 14 cells from N = 4 mice), CD2 (*n* = 14 cells from N = 4 mice), LFA-1 (*n* = 10 cells from N = 2 mice), membrane (*n* = 10 cells from N = 2 mice). i) Relative frequency distribution of TCR (*n* = 22 cells from N = 6 mice), CD2 (*n* = 24 cells from N = 6 mice), and LFA-1 (*n* = 18 cells from N = 4 mice) patch distance ratios normalized to the microvillus length. Inset: patch distribution (%) with distance ratio <1 (below the tip) and ≥1 (at or beyond the tip). j-k) Relative frequency distribution of the distance ratios between (j) CD2 and TCR (*n* = 10 cells from N = 2 mice) and between (k) CD2 and LFA-1 (*n* = 10 cells from N = 2 mice). Patches with the greatest distance to cell body within each CD2⁺TCR⁺ and CD2⁺LFA-1⁺ microvillus were used for analysis. Inset: patch distribution (%) with distance ratio <1 (below the tip) and ≥1 (at or beyond the tip). l) Scatter plot of the individual distance ratios between CD2:TCR and CD2:LFA-1 from (j) and (k), respectively. m) Cell score to compare tip preference using the normalized distance of furthest patch from the cell body. Scoring metric and equation can be found in methods. Sample sizes: CD2:TCR (*n* = 12 cells from N = 2 mice), CD2:LFA-1 (*n* = 10 cells from N = 2 mice), TCR:LFA-1 (*n* = 12 cells from N = 2 mice). Scale bar = 1 μm. Scale bar in (b) and (d) = 200 nm.

CD2 and LFA-1 differ in both binding affinity and structural organization. CD2 functions as a costimulatory receptor and its cytoplasmic tail associates with signaling and adaptor proteins, including Lck, Fyn and CD2AP.^[30]^ At the IS, CD2 forms a CD2-rich compartment that amplifies TCR signaling.^[36]^ CD2 binds CD48 in mouse (CD58/LFA-3 in human) with relatively weak affinity *K_d∼_*90*μM* (*K_d∼_*100*μM* in human), a property thought to support dynamic, flexible scanning during early antigen recognition.^[30,37–39]^ In contrast, LFA-1 is an integrin that recognizes ICAM-1 and ICAM-2 on APCs. In resting T cells, most LFA-1 molecules exist in a bent, low-affinity conformation, whereas TCR activation rapidly induces LFA-1 transition to an extended, high-affinity conformation that supports strong adhesion and stabilizes the IS.^[40,41]^ Thus, while CD2 is closely associated with TCR signaling, LFA-1 primarily mediates firm adhesion. A second major distinction lies in ectodomain size, which constrains where these receptors can reside within the IS. Since the membrane separation at TCR-pMHC contacts is ∼15 nm, receptor-ligand pairs that are significantly longer are sterically excluded from these close-contact zones.^[15,23]^ CD2 has a short ectodomain (∼7.5 nm), yielding a CD2-CD48 bond length of ∼15 nm that is compatible with the TCR-pMHC close-contact region.^[38,42]^ In contrast, fully extended LFA-1-ICAM-1 pairs have a bond length of ∼40 nm, which segregates them away from the tight TCR-pMHC contacts (see model, **Figure 8**).^[43,44]^ These distinct functional and structural differences motivate comparing the spatial organization of CD2 and LFA-1 relative to TCR on microvilli.

Previous work has shown that CD2 is enriched on microvilli, whereas LFA-1 is more uniformly distributed on the cell body.^[24,45–47]^ Jenkins et al. demonstrated that microvilli help T cells penetrate the APC glycocalyx and highlighted CD2’s role in stabilizing close contacts during antigen recognition.^[46]^ These results suggest that CD2 and LFA-1 are pre-organized into distinct membrane domains prior to activation. To define this pre-organization, we first measured the patch density, size (diameter, area, volume, number of voxel), and sphericity for each receptor (**Figure 3o-p**, **Figure S4d-g**). Then we mapped the distribution of CD2, LFA-1, and TCR on T cell membrane and compared their nanoscale organization on microvilli (**Figure 4f-h**). LFA-1 showed the highest overall density on the membrane, and TCR, CD2, and LFA-1 each displayed 1.10±0.14 patches per microvillus, 1.24±0.21 patches per microvillus and 1.18±0.15 patches per microvillus, respectively (**Figure 3o**, **Figure 4e**).

We next asked how frequently individual microvilli contained each receptor and how extensively patches occupied microvillar volume. We found ∼78% of microvilli were positive for TCR and LFA-1, whereas ∼85% were CD2-positive (**Figure 4f**). CD2 overlapped ∼60% of the MV region volume (**Figure 4g**), compared to ∼40% of that with TCR and LFA-1. To quantify enrichment within microvilli relative to the whole cell membrane, we calculated the fraction of total cellular fluorescence intensity localized in the MV region. A greater fraction of CD2 intensity was localized to the MV region than TCR or LFA-1 (∼30% CD2 vs. ∼20% TCR and LFA-1), while TCR and LFA-1 showed distributions comparable to that of the membrane marker (**Figure 4h**). Because our 3D analysis excludes the base of microvilli and short protrusions, these fractions likely underestimate the true microvillar contribution. Using an independent 2D slice-based analysis that includes shorter microvilli and the entire microvillar length, we confirmed that CD2 exhibits microvillar enrichment above the membrane marker, whereas TCR and LFA-1 were comparable to the membrane marker (**Figure S5**). Together, these analyses suggest differential preference of receptors on microvilli, with CD2 displaying a stronger preference for microvillar localization than TCR and LFA-1.

We then examined where within individual microvilli these patches reside—tip, shaft, or near the cell body. For each patch, we computed a “distance ratio” by dividing the maximal distance of the patch from the cell body by the microvillus length (**Figure 4i**, **Figure S4h-i**). Patches with a distance ratio <1 indicate that the patch does not reach the maximal edge of the microvillar tip or the patch occupies the shaft or proximal region. Patches with distance ratio ≥1 indicate that the patch is located at the tip. Using this metric, 69% of TCR patches and 68% of LFA-1 patches had a distance ratio <1, most of them between 0.8 and 1, indicating they occupy the distal region of the microvilli near the tip. In contrast, 54% of CD2 patches had a distance ratio <1, implying that nearly half of CD2 patches are located at the microvillar tips (**Figure 4i**). To directly compare CD2 and TCR within the same microvilli, we computed the distance ratio for CD2 and TCR patches in microvilli containing both receptors and found that 71% of CD2 patches were closer to the tips than their paired TCR patches (**Figure 4j, 4l**). A similar pattern was observed for CD2 versus LFA-1, with 69% of CD2 localized further toward the tips than LFA-1 (**Figure 4k-l**).

To summarize these pairwise preferences, we devised a tip-preference score for receptor pairs (e.g., CD2:LFA-1). For each microvillus, we identified the furthest patch from the cell body for each receptor in the pair, measured its distance to the cell body, and subtracted the two distances (e.g., *d_CD_*_2_ − *d_LFA_*_−1_), normalizing by the microvillus length (**Figure 4i**). For microvilli containing only one of the receptors (e.g., only CD2, no LFA-1), the distance of the absent receptor was set to zero. The final score per cell was the average of these normalized differences.

Positive scores indicate that the first receptor in the pair is biased toward the tip relative to the second. For CD2:TCR and CD2:LFA-1, average scores were positive, consistent with CD2 preferentially localizing toward microvillar tips. By contrast, the average score of TCR:LFA-1 was near zero, indicating no clear tip bias between these two receptors (**Figure 4m**).

Collectively, these results show that, relative to the membrane marker, CD2 is preferentially enriched on microvilli, and that CD2 exhibits a stronger preference for microvillar tips with a more distal localization relative to TCR and LFA-1.

### CD2 exhibits higher colocalization with TCR than LFA-1, specifically in the microvilli region

We next examined the spatial association between CD2, LFA-1, and TCR both on and off microvilli (**Figure 5a-d, Figure S6a-d**). In OT-I T cells labeled with TCR and CD2, TCR and CD2 patches showed close association, as highlighted by the white arrows (**Figure 5a-b**). In contrast, in T cells labeled with TCR and LFA-1, although some TCR and LFA-1 patches overlapped, their colocalization was visibly less pronounced than that of TCR and CD2 (**Figure 5c-d**). Consistent with these qualitative observations, TCR exhibited greater overlap with CD2 than with LFA-1, with ∼65% and ∼50% of TCR overlapping CD2 and LFA-1, respectively (**Figure 5e**). This difference was further accentuated when we restricted the analysis to patches within the MV region: ∼80% of TCR in microvilli overlapped CD2, whereas only ∼45% overlapped LFA-1 (**Figure 5f**). By contrast, in the non-MV (cell body) region, TCR overlap with CD2 (∼65%) and LFA-1 (∼60%) did not show a significant difference (**Figure 5g**). When we quantified CD2 overlap with TCR and LFA-1, we observed similar levels of occupancy across the entire membrane (**Figure S6e-g**), with no statistically significant differences when the analysis was restricted to the MV region or non-MV region. Examination of LFA-1 overlap with CD2 and TCR revealed that LFA-1 colocalized more with CD2 than with TCR, indicating that TCR and LFA-1 show the weakest pairwise spatial association (**Figure S6h-j**). Together, these results show that TCR is more closely associated with CD2 than with LFA-1, both across the whole cell and within microvilli, and that TCR-CD2 association is significantly enhanced within microvilli relative to the whole cell (paired two-tailed t-test, **p = 0.0070; **Figure 5e-f**).

**Figure 5.**
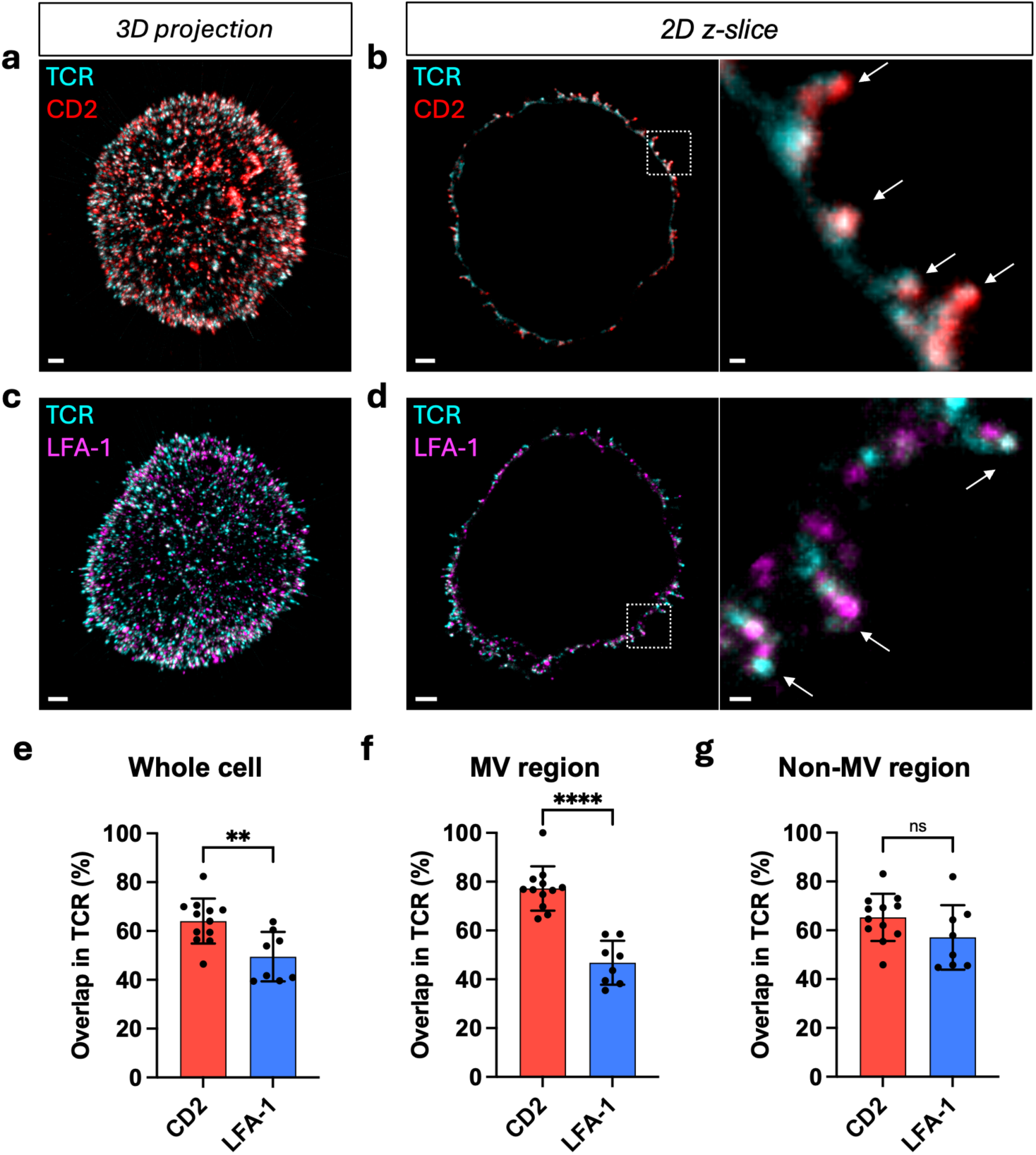
CD2 shows higher association with TCR patches in the MV region. a-b) Post-expansion images of OT-I T cell labeled with anti-TCR-AF488 (cyan) and anti-CD2-AF555 (red) in (a) 3D maximum projection and (b) 2D z-slice of the whole cell and magnified view of the region within the white box. c-d) Post-expansion image of OT-I T cell labeled with anti-TCR-AF488 (cyan) and anti-LFA-1-AF555 (magenta) in (c) 3D maximum projection and (d) 2D z-slice image of the whole cell and magnified view of the region within the white box. White arrows are pointing at the microvilli tips. e-g) Colocalization of CD2 and LFA-1 patches with TCR patches in (e) whole T cell membrane, (f) MV region, and (g) non-MV (cell body) region. In (e-g), data are presented as mean ± S.D. Statistical significance tested using unpaired two-tailed Welch’s t-test (whole cell: \*\**p* = 0.0055; MV region: \*\*\*\**p* < 0.0001; non-MV region: *p* = 0.1584; ns = not significant). In (e) and (f), TCR overlap with CD2 in the whole cell vs. the MV region was compared using a paired two-tailed t-test \*\**p* = 0.0070). Sample size in (e-g): CD2 (*n* = 12 cells from N = 3 mice), and LFA-1 (*n* = 8 cells from N = 2 mice). Scale bar in (a) and (c) = 1 μm. Scale bar in (b) and (d) = 200 nm.

A notable observation from the post-expansion images was the presence of large CD2 patches. These large CD2 patches were frequently detected, whereas patches of comparable size were not observed for other receptors, including TCR, LFA-1, and CD45. To address the possibility of immunostaining-specific artifacts, various monoclonal and polyclonal antibody clones of CD2 were tested, all yielding similar results. As shown in **Figure S7**, these CD2 patches were localized and associated with the microvilli. The underlying cause of these substantial accumulations remains unknown and further investigation is needed to determine their mechanism and significance in T cell signaling.

### Redistribution of Receptors in Microvilli After T Cell Activation

At late stages of T cell activation, TCRs accumulate in the cSMAC and are subsequently removed from the plasma membrane by internalization, ectocytosis, or trogocytosis.^[26,48–50]^ How these processes reshape the surface distribution of TCR and other receptors remains unclear.

Here, we sought to define receptor redistribution shortly after T cell activation. To examine how TCR, CD2, and LFA-1 nanoscale organization changes, we activated OT-I T cells with pMHC-coated magnetic beads and performed ExM imaging after 1 h and 2 h of stimulation (**Figure 6a, 6d, 6g, Figure S8a, S8c, S8e**). For microvillus-specific measurements, we restricted our analysis to patches located within the MV region. We first quantified receptor patch density and number of patches per microvillus in resting cells (0 h) and in cells stimulated for 1 h or 2 h. TCR patch density and number of patches per microvillus decreased significantly following activation (**Figure 6b-c**), whereas no significant changes were detected in CD2 and LFA-1 over the same time course (**Figure 6e-f, 6h-i**).

**Figure 6.**
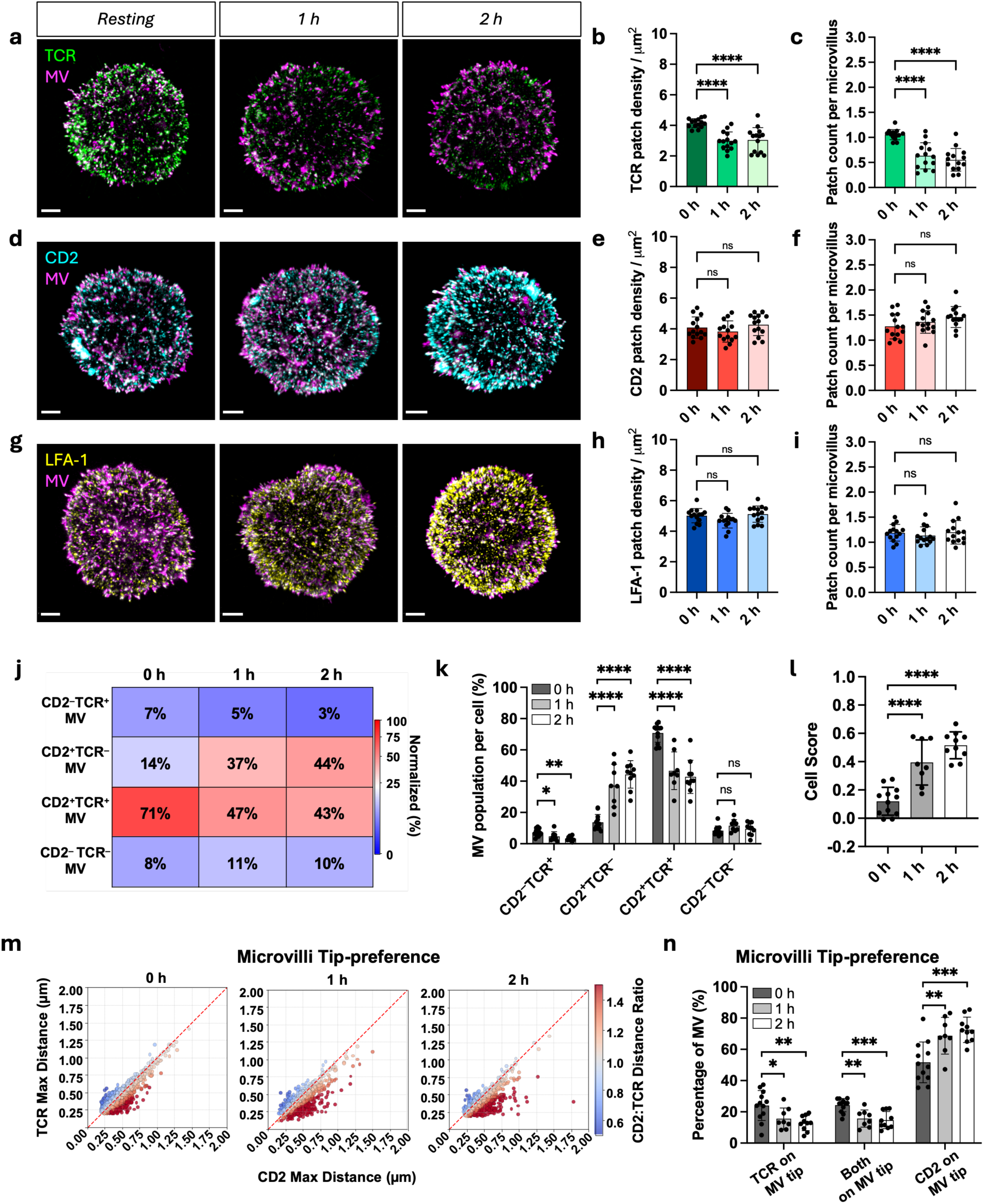
TCR patch density and occupancy in MV decreases while CD2 patch does not change post-activation. a) Post-expansion 3D image of OT-I T cell labeled with anti-TCR-AF488 (green) and WGA-CF633 (magenta) at resting (0 h), 1 h, and 2 h post-activation after co-incubation with pMHC-coated beads. b) Mean patch density of TCR per cell. c) Mean number of TCR patches localized per microvillus per cell. d) Post-expansion 3D image of a T cell labeled with anti-CD2-AF555 (cyan) and WGA-CF633 (magenta) at resting (0 h), 1 h, and 2 h post-activation after co-incubation with pMHC-coated beads. e) Mean patch density of CD2 per cell. f) Mean number of CD2 patches localized per microvillus per cell. g) Post-expansion 3D image of a T cell labeled with anti-LFA-1-AF555 (yellow) and WGA-CF633 (magenta) at resting (0 h), 1 h, and 2 h post-activation after co-incubation with pMHC-coated beads. h) Mean patch density of LFA-1. i) Mean number of LFA-1 patches localized per microvillus per cell. In (b-c), (e-f), and (h-i), data are presented as mean ± S.D. Statistical significance was determined using ordinary one-way ANOVA with Dunnett’s test against 0 h. Density in (b), (e), (h): (TCR 0 h vs. 1 h: \*\*\*\**p* < 0.0001; TCR 0 h vs. 2 h: \*\*\*\**p* < 0.0001; CD2 0 h vs. 1 h: *p* = 0.5306; CD2 0 h vs. 2 h: *p* = 0.7053; LFA-1 0 h vs. 1 h: *p* = 0.1582; LFA-1 0 h vs. 2 h: *p* = 0.8148; ns = not significant). Number of patches per microvillus in (c), (f), (i): (TCR 0 h vs. 1 h: \*\*\*\**p* < 0.0001; TCR 0 h vs. 2 h: \*\*\*\**p* < 0.0001; CD2 0 h vs. 1 h: *p* = 0.4936; CD2 0 h vs. 2 h: *p* = 0.0667; LFA-1 0 h vs. 1 h: *p* = 0.6730; LFA-1 0 h vs. 2 h: *p* = 0.9893; ns = not significant). j) Quadrant plot of the mean microvillar CD2:TCR populations in resting (0 h), 1 h and 2 h activation. MV category: CD2^+^TCR^−^, CD2^−^TCR^+^, CD2^+^TCR^+^, and CD2^−^ TCR^−^. Color gradient indicates change in microvillar population ranging from 0% (blue) to 100% (red). k) Bar graph showing fraction (%) of microvilli per cell in each CD2:TCR expression quadrant from (j). Statistical significance was determined using ordinary one-way ANOVA with Dunnett’s comparisons test compared against 0 h. (CD2^−^TCR^+^ 0 h vs. 1 h: \**p* = 0.0473; CD2^−^ TCR^+^ 0 h vs. 2 h: \*\**p* = 0.0012; CD2^+^TCR^−^ 0 h vs. 1 h: \*\*\*\**p* < 0.0001; CD2^+^TCR^−^ 0 h vs. 2 h: \*\*\*\**p* < 0.0001; CD2^+^TCR^+^ 0 h vs. 1 h: \*\*\*\**p* < 0.0001; CD2^+^TCR^+^ 0 h vs. 2 h: \*\*\*\**p* < 0.0001; CD2^−^TCR^−^ 0 h vs. 1 h: *p* = 0.1143; CD2^−^TCR^−^ 0 h vs. 2 h: *p* = 0.6728; ns = not significant). Sample sizes in (a-i): *n* = 14 cells from N = 3 mice for all conditions. l) Cell scores to compare the microvilli tip preference of TCR and CD2 in resting and activated T cells using the normalized distance of furthest patch from the cell body. Scoring system and equation can be found in methods. Data are presented as mean ± S.D. Statistical significance was determined using ordinary one-way ANOVA with Dunnett’s comparisons test against 0 h (0 h vs. 1 h: \*\*\*\**p* < 0.0001; 0 h vs. 2 h: \*\*\*\**p* < 0.0001). m) Scatter plot of the max distances of TCR (y-axis) and CD2 (x-axis) measured from the cell-body reference surface within each CD2⁺TCR⁺ microvillus at resting (0 h), 1 h and 2 h bead activation. Data points are colored by the distance ratio, with white indicating ratio = 1, blue indicating ratio < 1 and red indicating ratio > 1. n) Bar graph of the distance ratio categories from (m) with TCR on MV tip (ratio<1), Both on MV tip (ratio=1), and CD2 on MV tip (ratio>1). Data are presented as mean ± S.D. Statistical significance was determined using ordinary one-way ANOVA with Dunnett’s comparisons test compared against 0 h. TCR on MV tip (0 h vs. 1 h: \**p* = 0.0402, 0 h vs. 2 h: \*\**p* = 0.0034); Both on MV tip (0 h vs. 1 h: \*\**p* = 0.0017; 0 h vs. 2 h: \*\*\**p* = 0.0003); CD2 on MV tip (0 h vs. 1 h: \*\**p* = 0.0051; 0 h vs. 2 h: \*\*\**p* = 0.0004. Sample sizes in (j-n): 0 h (*n* = 12 cells from N = 3 mice), 1 h (*n* = 8 cells from N = 2 mice), and 2 h (*n* = 10 cells from N = 2 mice). Scale bar = 2 μm.

We next categorized microvilli into four classes: CD2^+^TCR^−^, CD2^−^TCR^+^ (single positive), CD2^+^TCR^+^ (double positive), and CD2^−^TCR^−^ (double negative) (**Figure 6j-k**). With 1 h and 2 h activation, the CD2^−^TCR^+^ and CD2^+^TCR^+^ microvillus populations decreased from 7% to 3% and from 71% to 43%, respectively, whereas the CD2^+^TCR^−^ single-positive microvillus population increased from 14% to 44%.

To exclude the possibility that these changes arose from gross remodeling of microvillus number, we compared microvilli density across resting (0 h), 1 h, and 2 h activated T cells and observed no significant differences (**Figure S8g**). The fraction of microvilli containing TCR significantly decreased after 1 h and 2 h activation, whereas the microvilli containing CD2 or LFA-1 remained unchanged (**Figure S8h**). We confirmed this by quantifying receptor overlap within microvilli: TCR overlap in microvilli declined significantly after 1 h and 2 h activation, while CD2 and LFA-1 overlap within microvilli did not change (**Figure S8i**). When we examined the fraction of receptor patches residing in the MV region relative to the total number of patches in the whole cell, the fraction of TCR significantly decreased, whereas the fractions of CD2 and LFA-1 did not exhibit significant changes over time (**Figure S8j**).

We next analyzed how activation alters the intramicrovillar positioning of receptor patches. We first examined the distribution of the receptor distance ratio, calculated as the maximal distance of each patch from the cell body divided by the microvillus length. Compared with resting cells, the fraction of TCR patches with a distance ratio ≥1 decreased from 31% at 0 h to 24% at 2 h (**Figure S8b**), whereas the fraction of CD2 patches with a distance ratio ≥1 slightly increased from 46% at 0 h to 53% at 2 h (**Figure S8d**). Meanwhile, the fraction of LFA-1 patches with a distance ratio ≥1 was similar across time points from 32% to 33% (**Figure S8f**). Since these pooled patch distributions are descriptive, we next compared the receptor positioning statistically using per-cell metrics.

Activation alters both receptor occupancy in each microvillus (**Figure 6j-k**) and the position of receptors within each microvillus. To quantify both effects, we used the tip-preference scoring metric and compared the tip preference of TCR and CD2 across all microvilli of a cell, based on the distance of the furthest patch from the cell body normalized by microvillus length. Relative to resting cells (0 h), the CD2:TCR tip-preference score increased by 3.3-fold and 4.3-fold after 1 h and 2 h of pMHC-bead stimulation, respectively, indicating a progressive bias of CD2 toward the tips relative to TCR (**Figure 6l**).

Since the scoring metric includes all microvilli irrespective of receptor occupancy, it cannot distinguish between TCR loss from microvilli and a change in the relative positions of the two receptors. To compare the relative positions of TCR and CD2 directly within the same microvilli and independently of receptor detection, we restricted the analysis to microvilli containing both CD2 and TCR (CD2^+^TCR^+^ double positive). Within these microvilli, the fraction in which TCR extended further toward the tip than CD2 (ratio <1) decreased from 24% to 13% after 2 h of stimulation, whereas the fraction in which CD2 extended further than TCR (ratio >1) increased from 52% to 73% (**Figure 6m-n**). The increase in the tip-preference cell score therefore reflects not only TCR loss from microvilli (**Figure 6j-k**, **Figure S8h**) but also a shift of TCR away from the tip relative to CD2 within the same microvilli. Overall, upon activation, TCR is selectively depleted from microvilli and shifts toward more proximal regions relative to the resting state, whereas CD2 maintains patch density and increased tip preference after 1 h and 2 h of bead activation.

To test whether these changes also occur when T cells engage target cells that present ligands for CD2 and LFA-1, we co-cultured effector OT-I T cells with MC38-OVA tumor cells for 1 h and 2 h (**Figure S9a-c**). Consistent with bead stimulation, TCR patch density and occupancy in the MV region decreased significantly after 1 h and 2 h of co-culture (**Figure S9d, g**). In contrast, CD2 and LFA-1 microvillar occupancy did not change significantly (**Figure S9h-i**). CD2 and LFA-1 patch densities changed modestly at 2 h, with a slight decrease for CD2 (4.1 to 3.7 patches μm^-2^) and an increase for LFA-1 (4.0 to 5.0 patches μm^-2^), possibly reflecting engagement of their ligands on the tumor cells (**Figure S9e-f**).

### T-DC immunological synapse features TCR and CD2 colocalize in multifocal synaptic regions while LFA-1 is excluded

We next sought to define how CD2, TCR, and LFA-1 are organized within the IS formed between T cells and dendritic cells (DCs). Building on the canonical “bull’s-eye” synapse model, Demetriou et al. showed that CD2 is enriched at the outer edge of mature synapses, forming a CD2 “corolla” that engages additional costimulatory receptors and amplifies TCR signaling.^[36,51]^

However, most of these structures were characterized in T cell-B cell conjugates or on supported lipid bilayers. In contrast, because dendritic cells have a highly dynamic actin cytoskeleton and extensive membrane processes, mature synapses between T cells and DCs often consist of multiple discrete contact sites rather than a single centralized contact zone.^[52–54]^ It remains unclear how TCR, CD2, and LFA-1 are distributed within these multifocal synapses.

To examine mature T cell:DC synapses, we conjugated OT-I T cells with antigen-bearing DCs for 10 min, and used NanoMAP to map CD2, TCR and LFA-1 on T cell:DC conjugates. As shown in **Figure 7a-b**, TCR and CD2 co-occupied the synaptic contact regions, highlighted by white circles. In contrast, although TCR and LFA-1 were both present at the interface, TCR patches localized to central synaptic zones while LFA-1 patches were positioned more peripherally along the outer edge of the contact (**Figure 7c-d**). Line profiles across these synapses revealed closely aligned intensity peaks for TCR and CD2, whereas LFA-1 peaks were shifted away from the synapse center and largely excluded from regions of maximal TCR signal (**Figure 7e-f**).

**Figure 7.**
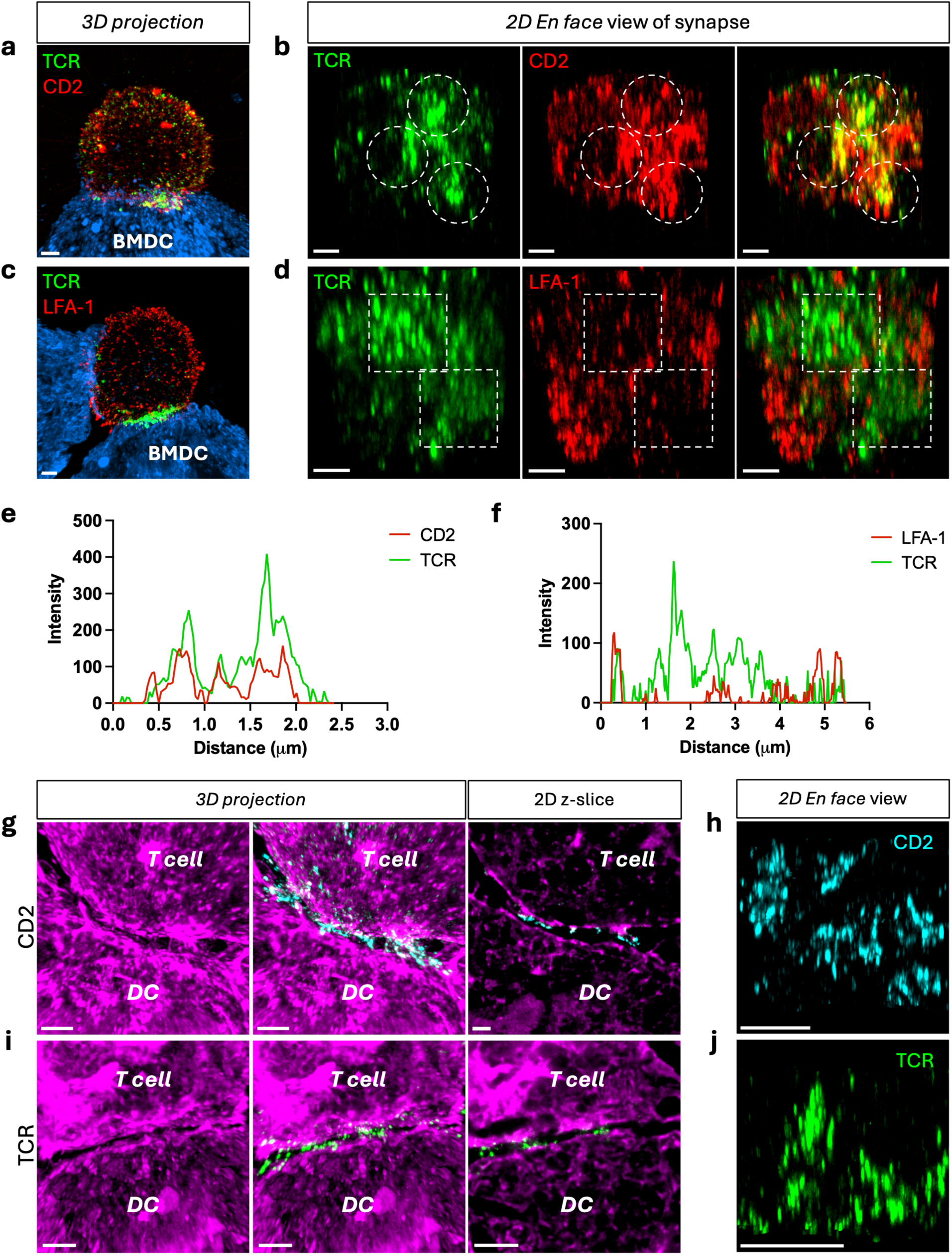
TCR and CD2 co-occupy multifocal tight contacts between T cell and dendritic cell. a) Post-expansion 3D image of OT-I T cell conjugated with a DC for 10 min. T cell is labeled with anti-TCR-AF488 (green) and anti-CD2-AF555 (red) and the DC membrane is labeled with WGA-CF633 (blue). b) *En face* 2D slice view of the synapse area between T cell: DC from (a). c) Post-expansion 3D image of OT-I T cell conjugated with a DC for 10 min. T cell is labeled with anti-TCR-AF488 (green) and anti-LFA-1-AF555 (red) and DC membrane is labeled with WGA-CF633 (blue). d) *En face* 2D slice view of the synapse area between T cell: DC from (c). e) Intensity plot of TCR and CD2 in the synapse *en face* slice view from (b). f) Intensity plot of TCR and LFA-1 in the synapse *en face* slice view from (d). Sample size in (e-f): intensity plots showing one representative region of interest (ROI) from *n* = 8 synapses across N = 2 mice. g) 3D maximum projection and 2D z-slice images with full-expansion in H_2O_ of a T cell: DC conjugate co-incubated for 10 min. T cell: DC is labeled with NHS-ester-Cy5 for pan-protein staining and T cell is labeled with anti-CD2-AF488 (cyan). h) *En face* 2D slice view of the contact area from (g). i) 3D maximum projection and 2D z-slice images with full-expansion in H_2O_ of a T cell: DC conjugate co-incubated for 10 min. T cell: DC is labeled with NHS-ester-Cy5 for pan-protein staining and T cell is labeled with anti-TCR-AF488 (green). j) *En face* 2D slice view of the contact area between T cell: DC from (i). Scale bar = 1 μm.

To resolve nanoscale membrane topology within the synaptic cleft, we used fully expanded samples in H_2O_ (7.3-fold expansion). In T cell:DC conjugates labeled with CD2 and NHS ester (pan-protein stain), CD2 patches were confined to the tightest contact regions and largely absent from adjacent loose-contact areas (**Figure 7g**). The *en face* view of the same synapses revealed multiple discrete CD2-enriched focal domains, with five prominent contact sites observed (**Figure 7h**). T cells labeled with TCR displayed a similar pattern: TCR patches were concentrated in tight contact zones and depleted from loose regions (**Figure 7i**), with four major TCR-enriched focal domains observed in the *en face* view (**Figure 7j**). Thus, within multifocal T cell:DC synapses, CD2 and TCR share a similar nanoscale distribution, occupying the tightest membrane appositions, whereas LFA-1 patches are segregated to more peripheral zones and excluded from these tight contact regions.

Although TCR and CD2 co-occupied synaptic contacts, their overall recruitment to the synapse exhibited different patterns. Whole-conjugate quantification showed that a larger fraction of TCR than CD2 localized to the synapse, whereas a larger fraction of CD2 remained in non-synaptic regions (**Figure S10a-e**). In addition, a larger fraction of TCR than CD2 was detected on the DC membrane, suggesting selective transfer of TCR (**Figure S10f-i**).

Our results show that TCR assembles into multiple discrete signaling domains that are co-occupied by CD2 but largely exclude LFA-1, a configuration that differs from the canonical bull’s-eye synapse with a CD2 corolla. Instead, our data are consistent with a multifocal synapse model and further demonstrate that the nanoscale organization of CD2 and LFA-1 is remodeled in this context. Given that Demetriou et al. implicated F-actin in CD2 corolla formation, an important future direction will be to determine how the dynamics and architecture of the actin cytoskeleton, and its coupling to CD2, differ across interactions with distinct APC types such as DCs and B cells.^[36]^

## Discussion

In this work, we established a platform, NanoMAP, that enables visualization and characterization of nanoscale membrane topology and receptor organization of T cell. We describe an experimental and computational workflow that combines ExM with 3D image analysis to profile receptors on defined membrane domains, such as microvilli. NanoMAP offers two expansion regimes: a 4.3-fold expansion optimized for mapping individual T cells, and a 7.3-fold full expansion optimized for resolving the IS.

NanoMAP offers several advantages over existing super-resolution approaches that have been used to study T cells and immune synapses. First, it provides volumetric 3D visualization of the entire T cell or T cell-APC conjugates, whereas techniques such as STORM/PALM are typically limited to 2D or a thin z-section of the cell.^[55]^ Second, it does not require specialized instrumentation such as electron microscopy or custom single-molecule setups; standard confocal microscopes are sufficient, yet the expanded samples still benefit from improvements in confocal resolution. Third, the labeling strategy is flexible and readily multiplexable. Because we rely on conventional antibodies rather than specialized fluorophores (as in STORM/PALM or STED), it is straightforward to map multiple targets: here we simultaneously visualized two receptor channels and one membrane channel, and additional targets can be included by choosing appropriate combinations of primary and secondary antibodies.^[28,29]^ The approach is also compatible with emerging multiplexed ExM development that can accommodate up to ∼10 targets in a single sample.^[27]^ Beyond surface receptors, the same workflow can be extended to intracellular molecules, such as Lck and cytoskeletal components (**Figure 2e**), making NanoMAP a versatile platform for profiling molecular organization within the T cell microvillar architecture. Beyond these imaging advantages, we report a panel of antibodies validated for ExM, together with a quantitative 3D segmentation workflow that maps receptors relative to microvilli across the entire cell surface.

NanoMAP is complementary to the cryo-expansion (cryo-ExM) platform recently reported by Lemaître et al., which integrates cryo-fixation with ∼4.5-fold expansion and mCLING membrane labeling for 3D T cells and immune synapses imaging.^[47]^ While cryo-fixation improves the preservation of actin networks and delicate membrane structures over standard PFA chemical fixation, cryo-ExM sample preparation requires cells to adhere to a coverslip. This substrate attachment can introduce mechanical stress and cell flattening. In contrast, NanoMAP processes T cells in suspension, which preserves native 3D cell morphology. Beyond sample preparation, NanoMAP delivers a quantitative image analysis pipeline for mapping receptors distribution relative to microvilli across the entire cell surface. Because the NanoMAP analysis framework processes 3D volumetric images, it is compatible with cryo-ExM images, combining improved structural preservation with quantitative receptor mapping.

NanoMAP also has limitations. First, although ExM improves effective resolution, it remains insufficient for accurately measuring molecular or cellular separations under 20 nm. Second, the profiling capability is constrained by the availability of antibodies that both recognize the target epitope and remain functional after the expansion and homogenization steps; this imposes a practical limitation on the receptor panel. Third, because imaging was performed with confocal microscopy, axial resolution is poorer than lateral resolution by approximately a factor of 2-3, which limits the effective resolution for synapses oriented perpendicular to the optical axis.^[28]^ Achieving near-isotropic resolution in all three dimensions will require integration with imaging modalities that provide improved z-resolution on expanded samples. Lastly, NanoMAP uses chemical fixation, which may imperfectly preserve fragile membrane structures compared with cryofixation.^[47]^ In addition, lipids are not retained after the harsh denaturation steps with ExM, therefore the membrane topology is visualized by glycoconjugates (WGA) or total protein (NHS ester) staining rather than by labeling the lipid bilayer. Since we identified microvilli by WGA staining, we validated the segmentation with β-actin labeling, which was detected in 94% of the segmented microvilli (**Figure S2d-f**).

We benchmarked NanoMAP’s mapping capability by reproducing key features of the T cell membrane previously resolved with other high-resolution methods. Specifically, we measured microvilli density and dimensions and obtained values that closely matched those reported by scanning electron microscopy^[33]^, demonstrating that NanoMAP captures microvillar architecture with sufficient fidelity. NanoMAP also recapitulated the differences in surface morphology between naïve and effector T cells, as well as the canonical bull’s-eye organization of the IS, consistent with prior studies.^[51,56,57]^

As part of this benchmarking, we mapped the distribution of CD45, a highly abundant T cell surface glycoprotein and phosphatase with a large ectodomain. On resting T cells, CD45 cytoplasmic tail dephosphorylates proximal TCR signaling components, thereby suppressing spontaneous TCR activation. As T cells form close contacts with APCs where TCRs engage pMHC, CD45 is sterically excluded from these contacts, facilitating the local phosphorylation required for productive TCR signaling.^[58–60]^ This segregation within the synapse was clearly resolved with NanoMAP.

Jung et al. previously used a distinct ExM workflow (30-40 nm resolution) and reported that CD45 on resting T cells is pre-excluded from microvillar tips, likely due to restricted diffusion of CD45 into ordered lipid domains at microvillar tips.^[61]^ In our 2D enrichment analysis (**Figure S5h**; ∼60 nm resolution), CD45 showed reduced microvillar-enrichment compared to the membrane marker WGA, suggesting partial pre-exclusion from microvilli. Visual examination of individual microvilli (both at ∼60 nm and ∼35 nm resolution) revealed heterogeneous degrees of pre-exclusion: some microvilli exhibited pronounced CD45 exclusion from the tip, others showed partial exclusion, and some displayed minimal pre-exclusion (**Figure S11**). ExM images from Jung et al. suggested more extensive pre-exclusion of CD45 from most microvillar tips, a pattern less frequently observed in our dataset. These differences may reflect higher labeling density in our samples or differences in imaging implementation (e.g., 4-fold expansion combined with Airyscan confocal in their study). Overall, our results indicate partial exclusion of CD45 from microvillar tips, which aligns more closely with findings from Kotowski et al.^[62]^, than with the extensive CD45 pre-exclusion reported for most microvilli tips. Because TCR on microvilli is distributed in proportion to membrane (**Figure 4h**, **Figure S5h**), this partial and heterogeneous pre-exclusion indicates that TCR and CD45 are partially segregated in resting T cells. This partial segregation complements rather than challenges the kinetic-segregation model.^[58]^ Jung et al. proposed that microvillar tips with CD45 depletion would reduce the amount of CD45 that needs to be excluded, potentially reducing the threshold for TCR triggering upon APC contact.^[61]^ Moreover, Kotowski et al. recently showed that partial exclusion is sufficient for TCR signaling.^[62]^ Therefore, partial pre-exclusion alone would not trigger signaling but could help reach the required level of exclusion. However, because CD45 partial exclusion occurs over ∼15 nm and below, which is beyond the effective resolution of NanoMAP, our data describe CD45 exclusion at the scale of individual microvilli and cannot resolve segregation at the molecular scale.

With NanoMAP, we profiled the effector OT-I T cell membrane and resolved the distribution of surface TCR, CD2, and LFA-1 relative to microvilli before, during, and after activation. Our data show that, prior to activation, CD2 patches are preferentially localized to microvilli, particularly at their tips, and enriched in microvilli relative to TCR and LFA-1. Within microvilli, TCR and CD2 patches display a high degree of spatial association, which suggests a positioning that could support efficient antigen surveillance. At mature synapses, CD2 remains closely associated with TCR within tight membrane contacts, whereas LFA-1 is segregated to more peripheral regions, where it can engage the APC membrane and provide high-affinity adhesion to stabilize the synapse (**Figure 8)**.

**Figure 8.**
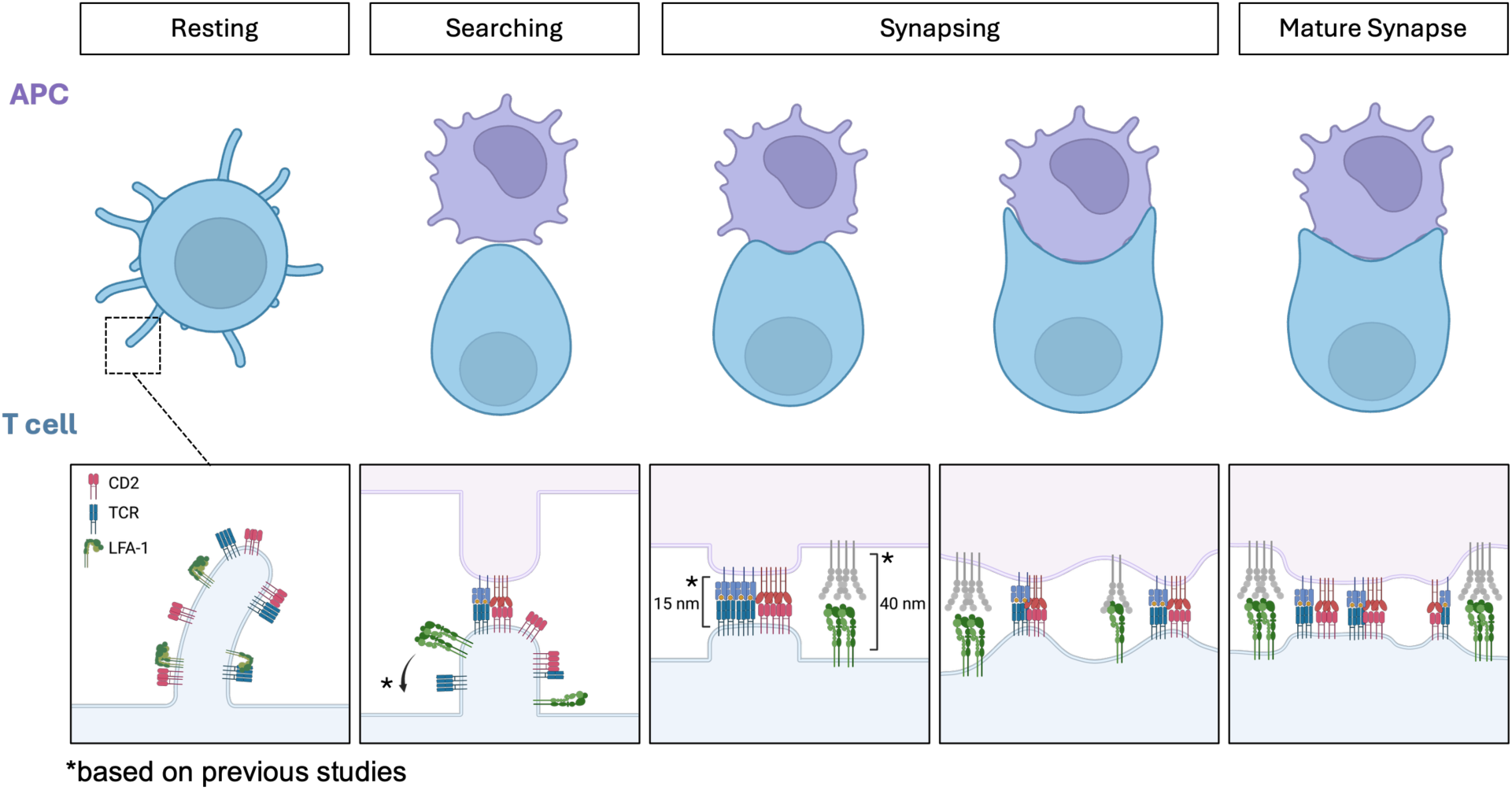
Model for CD2 localization at microvilli tips and its engagement with TCR during immunological synapse formation. In resting effector OT-I T cells, CD2 patches are enriched on microvilli relative to TCR and LFA-1, preferentially localized at their tips while exhibiting high spatial association with TCR patches. LFA-1 patches are localized on microvilli in the bent, low-affinity conformation prior to T cell activation. Upon antigen recognition at the searching stage, LFA-1 is expected to change to the extended (open) conformation as the T cell becomes activated.^[40,43]^ During synapse formation, TCR and CD2 co-occupy tight membrane contacts, consistent with the short bond lengths of TCR-pMHC and CD2-CD48 (∼15 nm),^[23,42]^ while LFA-1 is excluded from these contacts and localized in the peripheral regions, consistent with the longer binding length with ICAM-1 (∼40 nm).^[43,44]^ In mature synapses, TCR and CD2 remain closely associated, whereas LFA-1 is localized to the periphery. Elements marked with an asterisk (*), which depict LFA-1 conformation change and size-dependent segregation by bond length, are based on previous studies. ^[15,23,40,42–44]^

As T cells scan APC surfaces for antigen, microvillar tips act as initial contact points. Jenkins et al. reported that microvilli help T cells overcome the APC glycocalyx barrier and highlighted a role for CD2 in stabilizing close contacts during antigen recognition, while LFA-1 and CD45 are excluded from these sites.^[46]^ Our results extend this model by showing that microvilli are pre-organized in resting effector T cells with CD2 preferentially localized at microvillar tips, where it could help establish and stabilize nascent contacts, and CD2 and TCR are closely associated within microvilli, which could provide a nearby pool of TCRs for pMHC recognition. Although CD2 and TCR were previously reported to be enriched on microvilli, our work further resolves their spatial correlation within distinct microvillar subregions and outside microvilli, providing an architectural view of the pre-assembled microvillar signaling machinery.^[19,24]^ Our analyses in **Figure 4i** show that a large population of TCR patches are localized at the distal portions of microvilli, near but not reaching the tips, which aligns with previous reports of TCR residing at or near microvillar tips.^[24–26]^ In contrast, CD2 showed a larger population, relative to TCR and LFA-1, that more frequently extended to the tips (**Figure 4i**).

As part of this architecture, CD45 is partially excluded from microvillar tips, consistent with Kotowski et al. (**Figure S11**).^[62]^ LFA-1 showed a different pattern, with a distribution proportional to the membrane marker (**Figure 4h**, **Figure S5h**). This agrees with findings by Ghosh et al., who reported that most LFA-1 resides on the cell body, with ∼20% on microvilli and ∼5% at microvillar tips.^[45]^ They further show that LFA-1 on microvilli tips lies in close proximity to the chemokine receptor CCR7, enabling rapid chemokine-induced LFA-1 activation and ICAM-1 binding, thereby promoting T-cell arrest on chemokine-expressing surfaces.^[45]^

An important question raised by our findings is how differential receptor enrichment on microvilli is mechanistically controlled. Microvillar tips are high-curvature membrane regions and are therefore expected to be enriched in lipids and proteins that favor or stabilize curvature. Ordered lipid domains are thought to reside at microvillus tips; Jung et al. showed that such domains restrict CD45 diffusion, thereby contributing to CD45 exclusion from tips.^[61,63,64]^ Conversely, partitioning into these ordered domains may promote enrichment of specific receptors. Supporting this idea, Franke et al. demonstrated that palmitoylation is critical for CD4 accumulation at microvillar tips.^[65]^ Tetraspanins, a family of transmembrane proteins that favor high membrane curvature, are found at microvillus tips and can laterally organize partner receptors, providing another mechanism for selective enrichment.^[66–68]^ In addition, microvilli contain actin bundles, so interactions with the actin cytoskeleton offer yet another axis of control. For example, CD2 can interact with CD2AP (CD2-associated protein) and CIN85, which in turn bind actin, suggesting that actin-linked scaffolds may stabilize CD2 at specific microvillar locations.^[69,70]^ It is likely that a complex network of lipid partitioning, tetraspanin scaffolding, and cytoskeletal coupling cooperatively shapes receptor partitioning along microvilli. Therefore, a complete molecular architectural map of the microvillar machinery will be required to fully dissect these interconnected mechanisms.

At late stages of activation, T cells down-modulate surface TCR levels by internalization, thereby terminate signaling, and facilitate synapse disassembly.^[49,71]^ Recent studies further show that TCRs can be released from T cells to target cells across the synapse through ectocytosis, or trogocytic molting from microvilli.^[48,72]^ Park et al. reported that T cells activated on immobilized ligands shed microvilli as particles carrying multiple microvilli-associated proteins, including CD4, CD8, and CD62L, with surface TCR and microvilli recovering over 48 h.^[72]^ In **Figure 6**, we examined activation-induced alterations in T cell surface receptor organization. Following activation, TCR density and microvillar occupancy significantly decreased, whereas CD2 density remained unchanged. This activation-induced selective TCR depletion from the T cell surface is consistent with TCR downmodulation via internalization and ectocytosis from the synaptic center.^[48,49]^ However, because CD2 and TCR closely co-localize in microvilli (**Figure 5**), the selective loss of TCR from microvilli seems at odds with trogocytic molting, which should theoretically result in a comparable, or even greater, loss of CD2 relative to TCR. Notably, TCR and CD2 showed distinct recruitment patterns during synapse formation: TCRs underwent massive global recruitment to the synaptic region, whereas CD2 recruitment was localized and regional. Consequently, microvillar receptor composition undergoes extensive remodeling upon activation. Because synaptic microvilli become enriched in TCR relative to CD2, trogocytic molting occurring at the IS would naturally lead to predominant TCR transfer to the APC, as demonstrated in **Figure S10f-g**. Although our study evaluated membrane transfer after only 10 min of T cell:DC synapse formation, a time window insufficient to cover the extended molting timeframe reported by Park et al., our findings highlight that microvillar receptor organization is dynamically remodeled upon activation, setting synaptic microvilli apart from their resting state. Which specific structural and molecular components are remodeled in microvilli during activation, whether this reorganization is required to prime microvilli for trogocytic molting, and how distinct mechanisms differentially regulate receptor internalization, ectocytosis, and trogocytic molting upon activation remain to be elucidated.

It is important to note that we used pMHC-coated beads for T cell activation, which provide TCR stimulation but lack physiological ligands for CD2 and LFA-1. To examine receptor organization under more physiological conditions, we also characterized T cells after interaction with MC38-OVA tumor cells (**Figure S9**) and DCs (**Figure S10**). Additionally, the three receptors are not equivalent reporters of activation. While TCR is downmodulated upon signaling, LFA-1 is regulated through conformational changes, and CD2 is maintained or increased on activated T cells.^[31,40,71,73]^ Our results characterize the spatial distribution of the receptors rather than their functional states. Specifically, it was not verified whether the anti-CD11a clone used for LFA-1 labeling can distinguish open and bent conformations. Therefore, our conclusions are limited to the 1 h and 2 h activation time points examined in this report, and membrane organization beyond this time window cannot be inferred from our results. Pathways of TCR down-modulation may nonetheless differ across T cell subsets (CD8 vs CD4) and receptor-ligand combinations, potentially shifting the relative contributions of internalization, ectocytosis and trogocytic molting.

Our quantitative analyses and conclusions were drawn from effector OT-I CD8^+^ T cells, which allow antigen-specific activation under controlled conditions. NanoMAP is also applicable to human T cells as shown in **Figure S1i**. We visually observed CD2 at microvillar tips in primary human CD8^+^ T cells (**Figure S1i**; not quantified). However, quantitative findings of receptor organization in mouse CD4^+^ T cells and human T cells remain to be determined and will require further study.

In conclusion, our study leverages ExM-based 3D imaging and nanoscale profiling to explore how TCRs, adhesion receptors, and microvillar architecture are coordinated across distinct stages of T cell activation. The data support a model in which, in effector OT-I T cells, receptors are organized on microvilli according to their biophysical properties and functional roles in a manner that could support antigen sensitivity and signal transduction. These findings highlight the need to define the full molecular architecture of the T cell microvillus and to uncover the mechanisms that regulate this nanoscale machinery.

## Methods

### Mice

C57BL/6-Tg (TcraTcrb)1100Mjb/J (OT-I) and C57BL/6 mice were either purchased from Jackson Laboratories (Bar Harbor, ME, USA) or bred in house. Animal maintenance and procedures were conducted according to a protocol approved by the Institutional Animal Care and Use Committee (IACUC) at the Carnegie Mellon University.

### Cell Culture

#### Cell Line

Jurkat cells (ATCC) were maintained in RPMI supplemented with 10% fetal bovine serum, 100 U mL^-1^ penicillin, 0.1 mg mL^-1^ streptomycin and 2 mM L-glutamine. MC38-OVA cells (a gift from Dr. Robert Eil, OHSU) were maintained in DMEM supplemented with 10% fetal bovine serum, 100 U mL^-1^ penicillin, 0.1 mg mL^-1^ streptomycin and 2 mM L-glutamine. Cells were cultured in an incubator at 37 °C with 5% CO_2._ Cells were passaged every 48 h-72 h.

#### Effector OT-I CD8^+^ T cell

OT-I T cells from OT-I transgenic mice were cultured in complete RPMI (RPMI supplemented with 10% fetal bovine serum, 100 U mL^-1^ penicillin, 0.1 mg mL^-1^ streptomycin, 2 mM L-glutamine, 10 mM HEPES, and 50 μM β-mercaptoethanol). Single-cell suspensions were prepared from the lymph nodes and spleens of 6-10-week-old OT-I transgenic mice. Splenocytes were incubated with 100 ng mL^-1^ SIINFEKL peptide in complete RPMI for 30 min at 37 °C and then washed three times. Cell suspensions from lymph nodes and splenocytes were mixed in a 1:1 ratio at a density of 2×10^6^ cells per mL in complete RPMI (25 mL) in a T-75 flask and kept in a tissue incubator at 37 °C with 5% CO_2._ After 48 h (day 2), additional complete RPMI (25 mL) was added to the flask and supplemented with 10 U mL^-1^ IL-2. T cells were cultured for an additional 48-96 h before use, with fresh complete RPMI and IL-2 added every 48 h.

#### Effector CD4^+^ T cells from B6 mice

CD4^+^ T cells were prepared from the lymph nodes and spleens of 6-10-week-old C57BL/6 mice and isolated using the EasySep Mouse CD4^+^ T Cell Isolation Kit (STEMCELL). Isolated cells were activated with Dynabeads Mouse T-Activator CD3/CD28 (Thermo Fisher) in complete RPMI supplemented with 30 U mL^-1^ IL-2. After 48-72 h (day 2-3), fresh complete RPMI with IL-2 was added and T cells were cultured for an additional 48 h before use, with fresh complete RPMI and IL-2 added every 48 h.

#### Human CD8^+^ T cells

Human primary CD8^+^ T cells derived from human peripheral blood mononuclear cells (PBMC) were isolated using the EasySep Human CD8^+^ T Cell Isolation Kit (STEMCELL). After isolation, T cells were cultured in ImmunoCult-XF T cell Expansion Medium (STEMCELL) with 30 U mL^-1^ IL-2 and ImmunoCult Human CD3/CD28 T Cell Activator (STEMCELL) for activation and expansion. T cells were passaged every 48 h-72 h.

#### Bone marrow-derived Dendritic cell (DC)

DCs were prepared from C57BL/6 mice. The femur and tibia were isolated, and bone marrow was flushed using a needle and then mechanically dissociated to obtain a single-cell suspension. The cells were incubated in red blood cell lysis buffer for 5 min at room temperature, then washed and passed through 70 μm and 40 μm cell strainers. Cells were plated at a density of 1.2-1.5 x 10^6^ cells per mL in complete RPMI containing 20 ng mL^-1^ GM-CSF and incubated at 37 °C with 5% CO_2._ On day 3 and day 5, cultures were replenished with fresh media to maintain a density of around 1.2 x 10^6^ cells per mL. Cell culture was supplemented with 20 ng mL^-1^ IL-4 on day 5 to promote dendritic cell differentiation. On day 6 or 7, nonadherent cells in suspension were collected and replated at 1.2-1.5 x 10^6^ cells per mL in the same IL-4-supplemented media and stimulated with 1 μg mL^-1^ lipopolysaccharide (LPS) for 5 h to overnight before experiments to induce DC maturation.

#### Activated T cell / post-activation

Peptide-MHC-functionalized beads (pMHC) were first prepared. Dynabeads M-280 Streptavidin (Thermo Fisher) were first washed in 1% BSA in DPBS and resuspended in DPBS (BSA-free). Biotinylated H-2K^b^–OVA^257-264^ (SIINFEKL) monomer (20 µg; NIH Tetramer Core Facility) was added to the bead suspension, and the mixture was incubated on a shaker at room temperature for 30 min. During the 30 min incubation, the mixture was gently mixed every 10 min to ensure uniform coating. After incubation, the mixture was washed 4-5 times with excess 0.1% BSA in DPBS to remove unbound protein and resuspended in 0.1% BSA in DPBS for subsequent use. To prepare activated T cells, live effector OT-I T cells were first isolated using Ficoll-Paque and washed with complete RPMI via centrifugation. For 1:1 cell-bead ratio, T cells at 1 × 10^6^ cells per mL were mixed with 25 μL mL^-1^ of the pMHC bead solution after it was pre-washed with sterile DPBS. T cells were seeded in a 6-well plate (1 mL per well) with pre-washed beads. The cell-bead mixture was incubated at 37 °C for 1-2 h (as required by the experiment). To remove the beads, the mixture was transferred to a tube and placed on an EasySep magnet to harvest the activated T cells.

#### T cell: MC38-OVA co-incubation

MC38-OVA cells were seeded in a 100 mm cell culture dish and cultured at 37 °C overnight to reach 80-100% confluency. Live effector OT-I T cells were isolated using Ficoll-Paque and washed with complete RPMI before being added to the culture dish at a T cell-to-tumor cell ratio of 2:1, carefully without disrupting the adherent cells. Cells were co-incubated at 37 °C for 1 h or 2 h. Non-adherent T cells were collected and ready for subsequent ExM preparation.

#### Generation of naïve T cells

Naïve T cells were extracted from the lymph nodes of OT-I-TCR transgenic mice. Naïve OT-I T cells were then isolated using the EasySep Mouse CD8^+^ T Cell Isolation Kit (STEMCELL) and underwent ExM sample prep immediately following isolation.

### Preparation of T cell for ExM

#### Effector T cells

From the effector OT-I T cell culture flask, live cells were isolated using Ficoll-Paque and washed by centrifugation in complete RPMI. T cells were resuspended in complete RPMI and stained with primary antibodies at a concentration of 5 μg mL^-1^ for 30 min on ice. After rinsing, cells were fixed for 30 min on ice in fixation buffer (4% paraformaldehyde (PFA) and 0.01% glutaraldehyde in 1x PBS). Fixed cells were thoroughly washed with 1x PBS. Cells labeled with primary antibodies were resuspended in 1% BSA and stained with secondary antibodies at a concentration of 2 μg mL^-1^ for 1 h at room temperature. Cells were then washed with 1x PBS and ready to be gelled for ExM. It is important to note that both staining and fixation are performed on ice to limit crosslinking. Although this live-labeling approach has been validated in previous work,^[24]^ these measures do not fully exclude the possibility of antibody-mediated crosslinking.

#### Conjugates

To enable IS formation, effector OT-I T cells were incubated with antigen-pulsed DCs (100 ng mL^-1^ SIINFEKL peptide, 30 min at 37°C). Mature DCs (day 7, LPS-treated) were harvested by removing the media, then adding fresh complete RPMI to the flask and detaching adherent cells with a cell scraper. Harvested DCs were washed 3 times in complete RPMI by centrifugation. Dead cells were removed from T cell culture using Ficoll-Paque. DCs and T cells were then mixed in an Eppendorf tube at a 1:1 ratio at a total cell density of 1×10^7^ cells per mL in complete RPMI. The T cell-DC mixture was centrifuged at low speed (300 × g) for 1 min and incubated at 37°C. Incubation time varied depending on the stage of synapse formation for visualization. After incubation, cells were plated onto poly-D-lysine-coated coverslips. Excess cells floating in suspension were rinsed, and the cells on the coverslip were fixed in fixation buffer (4% PFA and 0.01% glutaraldehyde in 1x PBS) for 15 min at room temperature. The cells were washed thoroughly with 1x PBS to remove the fixation buffer, then stained with primary antibodies at 5 μg mL^-1^ in 1% BSA for 1 h at room temperature. For labeling intracellular proteins, the cells were first permeabilized with 0.05% Triton X-100 for 10 min, then labeled with primary antibodies. After washing the cells three times, they were stained with secondary antibodies at 2 μg mL^-1^ in 1% BSA for 1 h at room temperature. The T cell-DC conjugates were washed with 1x PBS and were ready to be gelled for ExM.

### Preparation of T cells on Supported Lipid Bilayer (SLB)

#### SLB preparation

Phospholipid mixtures in chloroform were prepared with 96.5% 1-palmitoyl-2-oleoyl-sn-glycero-3-phosphocholine, 2% 1,2-dioleoyl-sn-glycero-3-[(N-(5-amino-1-carboxypentyl)iminodiacetic acid)succinyl] (nickel salt), 1% 1,2-dioleoyl-sn-glycero-3-phosphoethanolamine-N-(cap biotinyl) (sodium salt) and 0.5% 1,2-dioleoyl-sn-glycero-3-phosphoethanolamine-N-[methoxy(polyethylene glycol)-5000] (ammonium salt). Lipids were mixed in a round-bottom glass flask, dried first under nitrogen, and then overnight in vacuum. Crude liposomes were generated by rehydrating the dried lipid film in 1x PBS to a final total lipid concentration of 4 mM and incubating for 1 h at room temperature. Small unilamellar liposomes were produced by extrusion through 100-nm Track-Etch membranes using a LiposoFast extruder. Four-well Nunc Lab-Tek II chambered cover glasses were cleaned by immersion in 5% Hellmanex at 55 °C, held at room temperature overnight, and then extensively rinsed with 18 MΩ water and air-dried. Glass coverslips were cut to fit each well and placed at the bottom of the chamber. The inserted coverslips were cleaned with 3 M NaOH (250 μL) at 55 °C for 15 min, rinsed extensively with 18 MΩ water, and the NaOH treatment was repeated. After a final extensive rinse with 18 MΩ water, chambers were dried under compressed air. To form SLBs, the liposome solution was added onto the inserted coverslips and incubated for 30 min at room temperature. To remove excess liposomes, each well was rinsed with 1x PBS (16 mL) using repeated additions of 1x PBS (0.5 mL) and aspiration to avoid drying out the SLB. Bilayers were blocked with 2% BSA in 1x PBS for 30 min at room temperature. Streptavidin (50 ng per well) was added for 30 min, followed by 1x PBS rinsing. Protein mixtures containing ICAM-1 with his-tag (126 ng) and biotinylated H-2K^b^–OVA^257-264^ (SIINFEKL) monomer (12 ng) in 2% BSA were added to the bilayers and incubated for 30 min to functionalize the SLBs. After functionalization, wells were rinsed with 1x PBS. Imaging medium which consisted of phenol red-free complete RPMI was added to the wells. Functionalized SLBs were equilibrated at 37 °C in a humidified incubator before adding cells.

#### Synapse formation on SLB

Effector OT-I T cells were added to SLB-functionalized coverslips and incubated with the bilayers for 7.5 min at 37 °C to form immune synapses. Samples were then fixed with 4% PFA and processed for immunostaining. Fixed cells were incubated with 5 μg mL^-1^ primary antibodies in 1% BSA for 1 h at room temperature, followed by 2 μg mL^-1^ fluorophore-conjugated secondary antibodies in 1% BSA for 1 h at room temperature. After completion of antibody staining, the coverslips were carefully removed from the chamber wells and used for subsequent fix cell imaging and/or expansion microscopy processing.

## Expansion Microscopy

### ExM method

The ExM protocol is based on the work of Klimas et al.^[29]^. The monomer solution was prepared with sodium acrylate, acrylamide, N, N-dimethylacrylamide, sodium chloride, N, N′-Methylenebisacrylamide, 10x PBS, and ddH_2O_. The gelling solution consisted of the monomer solution, N, N, N′, N′Tetramethylethylenediamine, methacrolein, and 2-Hydroxy-4′-(2-hydroxyethoxy)-2-methylpropiophenone.

For gelling suspension cells, the cells were centrifuged and resuspended in gelling solution (approximately 100 μL). The resuspended cells were dropped over a glass slide with a coverslip placed on top. To prevent the cells from flattening, two coverslip pieces can be placed at each end to create a chamber. The gel was polymerized under a UV (365 nm) curing lamp for 6 min. After the gel was polymerized, it was cut into smaller pieces using a razor blade and stored in Eppendorf tubes.

For gelling cells adhered to a surface, excess buffer was first removed from the cells. On a glass slide, gelling solution (∼ 60 μL) was dropped, and the cover slip with the adhered cells was placed over the slide, with the cells facing down so they were between the slide and the cover glass. The gel was polymerized under UV light for 6 min. After the gel was polymerized, it was cut into smaller pieces using a razor blade and stored in Eppendorf tubes.

Homogenization buffer containing urea, glycine, 0.5 M EDTA, tris base, and 10x PBS was pre-made.^[29]^ In the Eppendorf tubes, the gels were submerged in homogenization buffer and incubated for 1 h at 70 °C. After incubation, the gels were washed three times with wash buffer containing 0.1% C12E10 in 1x PBS.

To enhance the fluorescence signal, post-expansion staining was performed as previously described: the gels were stained with 5 μg mL^-1^primary antibodies in 5x SSC containing 0.2% Tween-20 overnight at room temperature.^[29,32]^ The gels were washed with 1x PBS three times. Lastly, the gels were stained with 2 μg mL^-1^ secondary antibodies in 5x SSC containing 0.2% Tween-20 for 2 h at room temperature and washed three times with 1x PBS. For cells that required membrane staining, the gels were stained with 5 μg mL^-1^ WGA or pan-protein labeling solution (NHS ester dye) in 1x PBS for 20 minutes at room temperature. The gels were washed with 1x PBS three times for at least 30 min before imaging.

### Full expansion (7.3-fold)

To achieve a higher expansion factor for enhanced resolution, full expansion can be achieved with higher water content. After completing antibody staining and/or membrane labeling, the gels were submerged in a 1:50 (1x PBS: H_2O_) solution and incubated for 10 min at room temperature. The solution was removed, and this step was repeated two more times. After the final solution was removed, the gels were ready for imaging.

### Confocal microscopy

Expanded samples were placed in a glass-bottom 6-well imaging plate and secured with plastic wrap to prevent the gel from drifting. Imaging was performed using a Nikon Eclipse Ti2 epifluorescence microscope equipped with a CSU-W1 spinning-disk confocal module and Hamamatsu camera. The system was controlled by NIS-Elements AR v.5.21.03 64-bit software. Images were taken using the following Nikon objectives: CFI Plan Apochromat VC ×60 C WI (1.2 NA). DAPI was excited with a 405 nm laser and imaged using a 450/50 emission filter; Alexa Fluor 488 with a 488 nm laser and 525/40 emission filter; Alexa Fluor 555 and DyLight 550 with a 561 nm laser and 607/36 emission filter; and Alexa Fluor 647 and Cy5 with a 640 nm laser and 685/40 emission filter.

### Expansion factor measurement with T cells (naïve/effector OT-I T cells, Jurkat cells)

T cells were labeled, fixed, and gelled as described above in *ExM Method*. Before the gels were cut, cells were imaged to obtain pre-expansion images. After homogenization, post-expansion staining, and expansion in 1x PBS or H_2O_, post-expansion images were acquired. For same-cell measurements, the cells imaged before expansion were located after expansion. The major axis diameter of the cell nuclei was measured in pre-expansion and post-expansion images, and the expansion factor was calculated as the ratio of the post-to pre-expansion measurements. For expansion factor comparison in separate cells, pre- and post-expansion images and measurements were obtained from separate cells. The expansion factor was calculated as the ratio of the mean diameter in post-expansion to pre-expansion, with 95% confidence intervals by bootstrap resampling (10,000 resamples).

### Isotropy and local distortion analysis

To quantify distortion introduced by expansion, we imaged the same field of view of OT-I T cells before and after expansion. Samples were labeled before expansion for pre-expansion imaging and labeled again after expansion, with all images acquired as z-stacks. The 2D distortion analysis described in the Magnify ExM method was used.^[29]^ For each ROI, the algorithm compared all z-slices of the two stacks, accounting for the expansion factor, and identified the best-matched pair of pre-expansion and post-expansion z-slices to ensure proper alignment for relative error measurements. From each pair of z-slices per channel, identifiable landmarks in both pre-expansion and post-expansion imaging were matched across the ROI for the distortion calculation.

The root-mean-square (RMS) length measurement error was then calculated as a function of measurement length, separately for each channel (**Figure S1e-f**). These values likely overestimate the true distortion. Because all channels were re-labeled after expansion, post-expansion staining can access epitopes and glycans that are inaccessible before expansion, and post-expansion imaging resolves features that are diffraction-limited in the pre-expansion image. As such, matching distinguishable features between pre-expansion and post-expansion images can be difficult given the post-expansion images often has increased number of labeled features with much higher resolution. Both scenarios create genuine differences between the two images that the algorithm can count as error.

### Expansion factor measurement with MC38 tumor cells

MC38 tumor cells were seeded on a 22 mm x 22 mm cover glass overnight. Before gelling, the cell membrane and nuclei were stained with 5 μg mL^-1^ WGA and 1 μg mL^-1^ DAPI, respectively. After washing the labeling solution, the adhered cells were gelled using the ExM method described above. Before cutting the gels, the cells embedded in the gelling solution were imaged for the pre-expansion images. After homogenization, the gels were re-stained with 5 μg mL^-1^ WGA and 1 μg mL^-1^ DAPI to enhance fluorescence. For post-ExM images, the identical cells from the pre-expansion images were located and obtained. The major axis diameter of the cell nuclei was manually measured in pre-expansion and post-expansion images. These measurements were used to calculate the linear ratio of the increase in size and determine the expansion factor (**Figure S1a-b**).

### Data Analysis

#### Image Preprocessing

Image files acquired by the confocal microscope were preprocessed in Imaris (Bitplane, Oxford Instruments). Preprocessing includes file format conversion and setting of the effective voxel size for expanded samples. Regions of interest (ROIs) were cropped to encompass the full 3D cell volume with minimal surrounding margin. The cropped volumes were then processed using the built-in background subtraction algorithm in Imaris to remove the background fluorescence (**Figure S2a-b**). The preprocessed files were saved in Imaris for further analysis.

#### Cell Body Detection and Segmentation

Cell bodies were segmented based on membrane staining using the cell detection algorithm in the Imaris for Cell Biologists package, and the resulting cell boundary were exported as surfaces. In cases where membrane staining was weak or discontinuous, standard detection performed poorly. To address this issue, two strategies were applied: (1) using the sum of normalized fluorescence intensities from all channels for cell segmentation instead of relying solely on membrane staining, and (2) enabling detection of segmented volumes followed by merging all segments to reconstruct the full cell volume.

#### Cell Region and Microvilli Region Segmentation and Masking

Distances from the cell surface were then calculated for each voxel within the ROI using the Imaris Distance Transformation function. The results were stored in a new “Distance to Cell” channel, where *r* = 0 marks the cell-body boundary, negative values indicate voxels inside the cell surface and positive values indicate voxels outside.

The “Cell Region” was defined as voxels within a distance range of -0.2 μm to 2 μm from the cell surface, excluding intracellular signals and fluorescence from neighboring cells (**Figure S2**). The “microvilli (MV) region” was defined as voxels within 0.15 μm to 2 μm from the cell surface to capture microvilli-associated signals while excluding the cell body membrane. The reference surface (*r* = 0) lies within the membrane fluorescence signal rather than at its outer edge. Because the membrane signal has an apparent thickness, its outer edge typically lies approximately 0.15 μm from the reference surface. Therefore, 0.15 μm was used as a spatial segmentation threshold to separate protruding membrane signal from the cell-body membrane signal. This threshold is an offset from the reference surface (*r* = 0) rather than an explicitly imposed minimum microvilli length. For each segmented microvillus, the length was defined as the length of the longest principal axis of an ellipsoid fitted to the segmented microvillus. Note that this segmentation method may exclude protrusions that do not extend beyond the threshold and may omit the proximal portion of protrusions that overlap with the cell-body membrane signal.

For each receptor and/or membrane channel, region-specific masks were applied by setting voxel values outside the defined regions to zero while retaining original values within the regions. This yielded three channels for each marker: a background-subtracted channel, a whole-cell region channel, and an MV region channel.

#### Receptor Patch and Microvilli Segmentation

Receptor patches and microvilli were segmented by generating surfaces using the built-in Imaris surface detection algorithm. The MV region channel of the membrane marker was used to identify microvilli. The whole-cell and/or MV region channels of receptors were used to identify receptor patches within the regions of interest. For quantitative analysis, binary masks were created from the detected surfaces, with voxel values set to 1 inside the surfaces and 0 outside.

#### Patch and Microvilli Colocalization Analysis

The colocalization of two different receptors, or a receptor patch with the microvilli, was measured by the extent of overlap between the two segmented patch volumes. The binary masks generated from the segmented patches were used for this analysis.

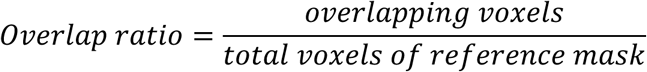

For example, to calculate the overlap of receptor A in receptor B, the number of voxels where the two masks overlap was divided by the total number of voxels in Receptor B

#### Nearby surface analysis

To quantify spatial relationships between individual receptor patches and microvilli surfaces, each segmented surface was masked and assigned a unique identifier using an in-house Imaris extension (SurfaceIndexMask, MATLAB). This tool generates a “Surface ID” channel for each marker in which all voxels within a given surface are labeled with the same unique ID. The corresponding 3D matrix of the surface index mask was also exported as a .mat file.

Using a second in-house Imaris extension (NearbySurface, MATLAB), secondary surfaces located within a defined distance of primary surfaces were identified. This analysis was used to detect receptor patches within a specified distance of each microvillus. Identified surfaces were extracted based on their unique IDs, and the results were exported in CSV format.

Next, the receptor patches were assigned to each microvillus based on voxel overlap. However, when a patch overlaps multiple microvillus segmentations, the patch is assigned to the microvillus with the greatest voxel overlap to avoid double counting. To examine localization of patches within the microvilli, the “distance ratio” was computed by dividing the maximal distance of each patch from the cell body by the length of its microvillus. The distance of each receptor patch was defined by its absolute distance-transform coordinate, measured from the cell-body reference surface (*r* = 0). To directly compare the distribution of two different receptor patches within the same microvilli (e.g. CD2 vs. TCR), the maximal distance of a CD2 patch was divided by the maximal distance of a TCR patch in the same microvilli.

#### Cell Scoring Analysis

Cell scoring analysis was performed to compare whether receptor A or receptor B preferentially localizes further toward the microvilli tip in each cell. The score was defined as:

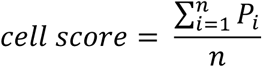

where *n* is the total number of microvilli of the cell, and *P_i_* is the score for the *i^t^*^ℎ^ microvillus. The microvillus score *P_i_* was calculated as:

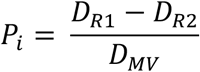

where *D_R_*_1_ and *D_R_*_2_ are the distances of the furthest patch of receptor A and receptor B on that microvillus, respectively, and *D_MV_* is the total length of the microvillus. This scoring scheme includes all microvilli, whether they contain both receptors, only one receptor, or neither receptor. For double-negative microvilli, both *D_R_*_1_ and *D_R_*_2_ are zero, yielding *P_i_* = 0, indicating no preference of either receptor for the tip. For microvilli containing only receptor A, *D_R_*_2_ = 0, resulting in a positive *P_i_* = *D_R_*_1/_*D_MV_*, which indicates a stronger preference of receptor A for the microvillus tip; conversely, microvilli containing only receptor B yield a negative *P_i_* = –*D_R_*_2/_ *D_MV_*, indicating a stronger preference of receptor B for the tip. The enrichment score for the cell is the mean of *P_i_* values for all microvilli.

#### Microvilli Enrichment Analysis

An in-house Python script (MV Enrichment Analysis) was used to quantify receptor enrichment in microvilli. The script takes multi-channel z-stack TIFF images exported from processed Imaris data files, including background-subtracted membrane and receptor channels, along with the “Distance to Cell” channel. Because microvilli at the top and bottom of the cell are mostly outside the imaging plane of a given z-slice, only z-slices from the mid-section of the cell were included for the analysis.

For each z-slice, all channels were transformed into polar coordinates, with the origin defined as the center of the cell at that z-slice; the x-axis encodes the angular position *Θ* and the y-axis encodes the radial distance *r* to the cell surface *r* = 0 on the cell boundary). Pixel intensities were normalized to generate angular intensity profiles in color scale, with the angular step size chosen to approximate the width of a microvillus (**Figure S5a-e**).

Microvilli were identified from the membrane intensity profile by detecting peak positions (distance to cell) at each angular step (**Figure S5f**). At each angular step, all peaks above a defined threshold were selected. A “global” peak position was defined as the mean position of all detected peaks, and the corresponding “global” peak width was defined as the distance between the left full width at half maximum (FWHM) boundary of the leftmost peak and the right FWHM boundary of the rightmost peak. Peak detection parameters were chosen to capture the full extent of each microvillus, including its base at the cell body. To account for membrane thickness, angular steps with a global peak position greater than 0.03 μm were classified as microvilli (*MV*), while the remaining angular steps were classified as cell body (*CB*). The global peak width was used to define the radial ranges of the corresponding *MV* and *CB* regions. This procedure was applied across all angles to define *MV* and *CB* regions for each z-slice (**Figure S5g**).

Receptor enrichment for each z-slice was calculated as the ratio *I_MV_*_/_( *I_MV_*_+_*I_CB_*_)_, where *I_MV_* _a_nd *I_CB_* _r_epresent receptor fluorescence intensity in the *MV* and *CB* regions, respectively. This value was normalized by the corresponding ratio calculated for the membrane channel in the same z-slice, and expressed as relative enrichment, (normalized ratio − 1) × 100% (**Figure S5g**). The normalized values were then averaged across all selected z-slices to obtain a per-cell enrichment metric. Positive enrichment values indicate preferential localization to microvilli, whereas negative values indicate exclusion, with 0 corresponding to the membrane baseline (**Figure S5h**).

#### Statistical Information

Normality was assessed before selecting the tests. For comparisons between two groups, we used an unpaired, two-tailed Welch’s t-test for measurements from different cells and a paired, two-tailed t-test for matched measurements from the same cells. For comparisons among three or more groups, we used ordinary one-way ANOVA followed by Tukey’s multiple-comparisons test when all pairwise comparisons were of interest, or Dunnett’s multiple-comparisons test when groups were compared with a single reference, such as the 0 h time point in activation experiments. GraphPad Prism 11 was used for statistical analyses and multiple comparisons. Statistical details of experiments and exact *P* values can be found in figure legends. Mean, standard deviation, and standard error of the mean were determined using GraphPad Prism 11.

To evaluate the sensitivity of our experimental design, we performed an effect-size sensitivity power analysis in Python (statsmodels, tt_ind_solve_power), calculating the minimum effect size (Cohen’s d) detectable with 80% power at a two-sided *a* of 0.05 for our typical sample sizes. Because our data were analyzed by one-way ANOVA followed by Tukey’s or Dunnett’s multiple comparisons, we used a Bonferroni-adjusted *a* for each pairwise comparison as a conservative approximation to these tests. For resting T cell experiments (typically ∼20 cells per condition), the minimum detectable effect size was d = 1.01 for tests with two comparisons (e.g., Dunnett’s test with three groups) and d = 1.06 for designs with three comparisons (e.g., Tukey’s test with three groups or Dunnett’s test with four groups). For bead-activation experiments (typically 14 cells per condition), the corresponding minimum detectable effect sizes were d = 1.23 and d = 1.30. For comparisons analyzed by Welch’s *t*-test, minimum detectable effect sizes were calculated using the actual group sizes and assuming equal variances in both groups with Python (tt_ind_solve_power), which closely approximates Welch’s *t*-test: d = 1.16 for **Figure 2d** (*n* = 10 and 17 cells), d = 1.35 for **Figure 5e-g** (*n* = 12 and 8 cells), d = 1.30 for **Figure S6e-g** (*n* = 12 and 9 cells), and d = 1.46 for **Figure S6h-j** (*n* = 9 and 8 cells).

### Materials

Detailed information on all materials is provided in Supporting Information, **Tables S1–S6**

## Acknowledgements

This work was supported by the Shurl and Kay Curci Foundation award to E.C. Y.Z. acknowledges funding from NIH grants RF1 MH129267, R01 EB035890, 1AY2AX000056, R01 CA301488, R.K. Mellon Foundation and the Eberly family professorship. E.L. acknowledges funding from NIH grant T32 GM133353 in the Interinstitutional Program in Cell and Molecular Biology.

We thank Dr. Robert Eil (Oregon Health & Science University) for kindly providing the MC38-ova cell line. Manual measurements of the microvilli were performed with UCSF ChimeraX, developed by the Resource for Biocomputing, Visualization, and Informatics at the University of California, San Francisco, with support from NIH R01-GM129325 and the Office of Cyber Infrastructure and Computational Biology, National Institute of Allergy and Infectious Diseases.

## Data Availability

Python code and in-house XTensions for Imaris used for data analysis in this paper are available on GitHub (https://github.com/Cai-Lab-CMU/NanoMAP-CD2). The repository also contains per-panel source data, and a table of antibodies tested for compatibility with NanoMAP. A representative dataset containing the Imaris files before and after analysis has been deposited in Zenodo (https://doi.org/10.5281/zenodo.23957253). Due to the large dataset and file size, additional data are available by request.

## Animal Ethics Statement

All experiments involving animals were approved by the Institutional Animal Care and Use Committee (IACUC) at Carnegie Mellon University and conducted in accordance with institutional guidelines.

## Declaration of Generative AI and AI-Assisted Technologies

Generative AI (Claude Sonnet 4.6, ChatGPT-5) was used to assist in drafting and debugging code for data analysis. All AI-assisted code was critically reviewed, revised, and validated by the authors before use. The authors take full responsibility for the final analysis code, its implementation, and the reported results.

During the preparation of this manuscript, generative AI (Gemini 2.5 Flash, Perplexity Sonar 2, Grammarly) was used to assist with language editing, grammar correction to improve the readability of the author-generated draft. Following the use of AI tools, the authors reviewed and revised the content as needed and take full responsibility for the accuracy, originality, and final content of the manuscript.

