## Supplementary Information for "Nanoscale 3D profiling of the T cell membrane reveals CD2 enrichment at microvilli tips, positioning adhesion near TCR zones in the immunological synapse"

**Supporting Information Summary**

1. Supplementary Figures and Figure Captions: **Figures S1–S9**
2. Movie Captions: **Movies S1–S3**
3. Supplementary Tables for Materials: **Tables S1–S6**

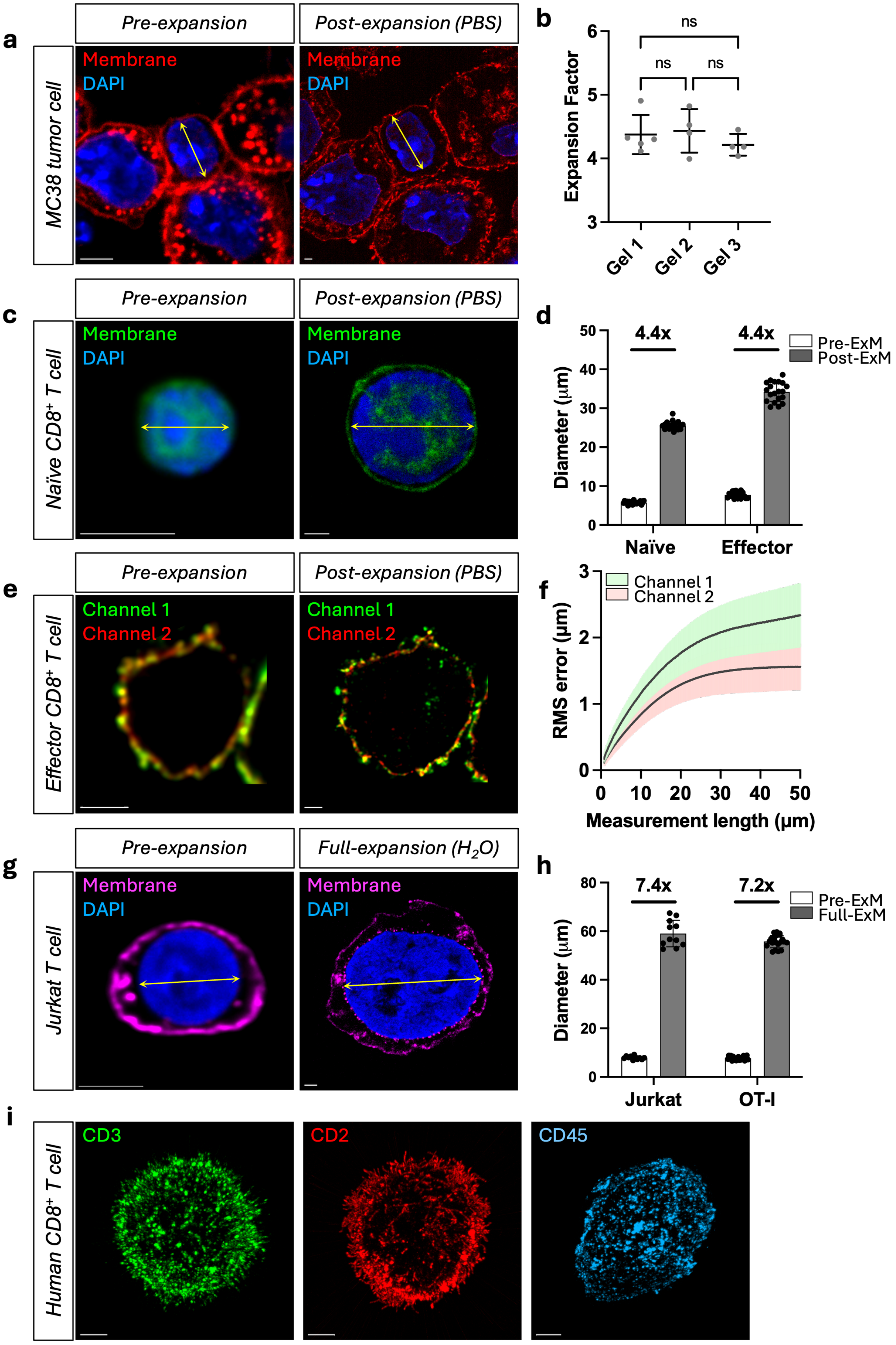
a
Pre-expansion
Post-expansion (PBS)
MC38 tumor cell
Membrane
DAPI
b
Expansion Factor
ns
Gel 1
Gel 2
Gel 3
c
Naïve CD8+ T cell
d
Diameter (µm)
4.4x
Pre-ExM
Post-ExM
Naïve
Effector
e
Effector CD8+ T cell
Channel 1
Channel 2
f
RMS error (µm)
Measurement length (µm)
g
Full-expansion (H2O)
Jurkat T cell
h
7.4x
7.2x
Full-ExM
Jurkat
OT-I
i
Human CD8+ T cell
CD3
CD2
CD45

**Figure S1. Calculation of expansion factor.**

a) Pre-expansion (left) and post-expansion images (right) in 2D z-slice of MC38 tumor cell labeled with WGA for membrane staining and DAPI. b) Expansion factor calculated using the major axis diameter of the nuclei as shown in a). Measurements were obtained from three separate gels and expanded independently. Sample sizes: gel 1 ($n$ = 5 cells), gel 2 ($n$ = 4 cells), gel 3 ($n$ = 4 cells). Data are presented as mean ± S.D. and statistical significance was determined using ordinary one-way ANOVA with Tukey's comparisons test ($p$ = 0.5489; gel 1 vs gel 2: $p$ = 0.9494; gel 1 vs gel 3: $p$ = 0.6902; gel 2 vs gel 3: $p$ = 0.5439; ns = not significant). c) 2D z-slice images of a naïve OT-I T cell labeled with WGA for membrane staining and DAPI in pre-expansion (left) and post-expansion (right) with 1x PBS. Yellow arrow = major axis diameter. d) Major axis diameter of naïve and effector OT-I T cell nuclei in pre-expansion and post-expansion in 1x PBS. Data are presented as mean ± S.D. Sample sizes: $n$ = 20 cells from N = 1 gel for naïve and effector OT-I T cell. e) Pre-expansion (left) and post-expansion images (right) in 2D z-slice of effector OT-I T cell with antibody-labeled receptor channel 1 and channel 2. f) Root-mean-square (RMS) measurement error versus measurement length (post-expansion scale) for antibody-labeled channels. Lines show the mean and shading shows ± SEM (C1: $n$ = 6, 2 gels; C2: $n$ = 8, 3 gels). g) 2D z-slice images of Jurkat cells labeled with WGA for membrane staining and DAPI in pre-expansion (left) and full-expansion with $H_2O$ (right). h) Major axis diameter of Jurkat cell and OT-I T cell nuclei in pre-expansion and post-expansion in $H_2O$. Data are presented as mean ± S.D. Sample sizes: Jurkat cell: $n$ = 11 cells, N = 1 gel; OT-I T cell: $n$ = 20 cells, N = 1 gel. In (d) and (h), the expansion factor was calculated as the ratio of the mean diameter in post-expansion to pre-expansion. i) Post-expansion images in 3D of human OT-I T cells isolated from peripheral blood mononuclear cells (PBMC) and labeled with anti-CD3, anti-CD2, and anti-CD45. Scale bar in (a), (c), (e), (g) = 5 μm. Scale bar in (i) = 2 μm.

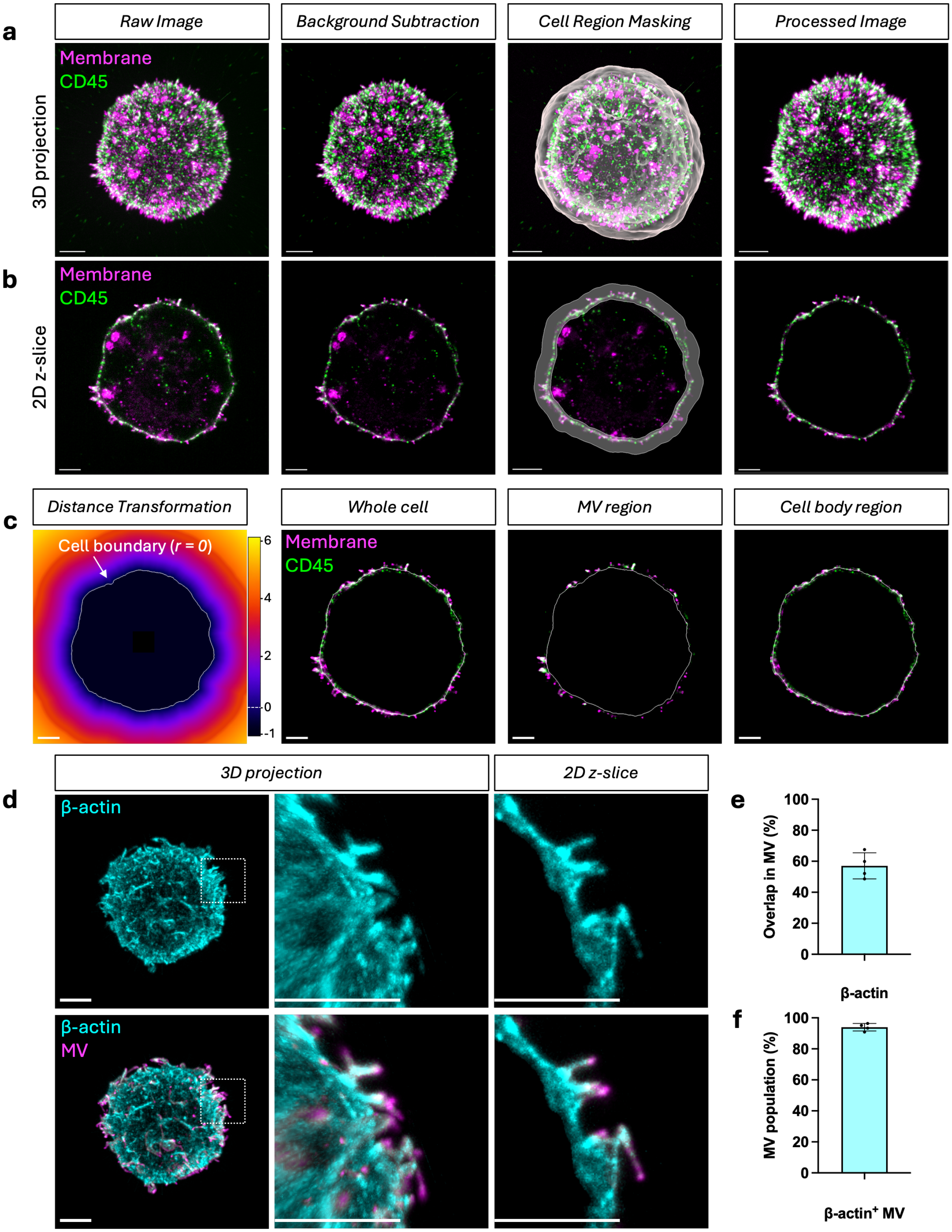
a
Raw Image
Background Subtraction
Cell Region Masking
Processed Image
3D projection
Membrane
CD45
b
2D z-slice
Membrane
CD45
c
Distance Transformation
Cell boundary (r = 0)
6
4
2
0
-1
Whole cell
MV region
Cell body region
Membrane
CD45
d
3D projection
2D z-slice
β-actin
β-actin
MV
e
Overlap in MV (%)
100
80
60
40
20
0
β-actin
f
MV population (%)
100
80
60
40
20
0
β-actin+ MV

**Figure S2. 3D image post-processing workflow and microvilli validation with β-actin**

a-b) Image post-processing steps of post-expansion images in (a) 3D maximum projection view and (b) 2D z-slice view. Representative image of OT-I T cell with anti-CD45 (green) and WGA-CF633 (magenta) staining. c) Distance transformation channel of a 2D z-slice with a cell boundary (white, $r = 0$) where voxels are assigned negative ($r < 0$) and positive ($r > 0$) distances. Images show fluorescence intensity of a whole cell, segmented MV region and segmented cell body (non-MV) region. Scale bar = 2 μm. d) Post-expansion images in 3D maximum projection view and 2D z-slice view of a OT-I T cell labeled with (top) β-actin and (bottom) β-actin with MV segmentation using WGA (magenta). Magnified image of the region within the white box. e) Mean occupancy (voxel overlap) of β-actin in the MV region per cell. f) Fraction of microvilli containing β-actin per cell. Sample sizes in (e-f): $n = 4$ cells from N = 1 mice.

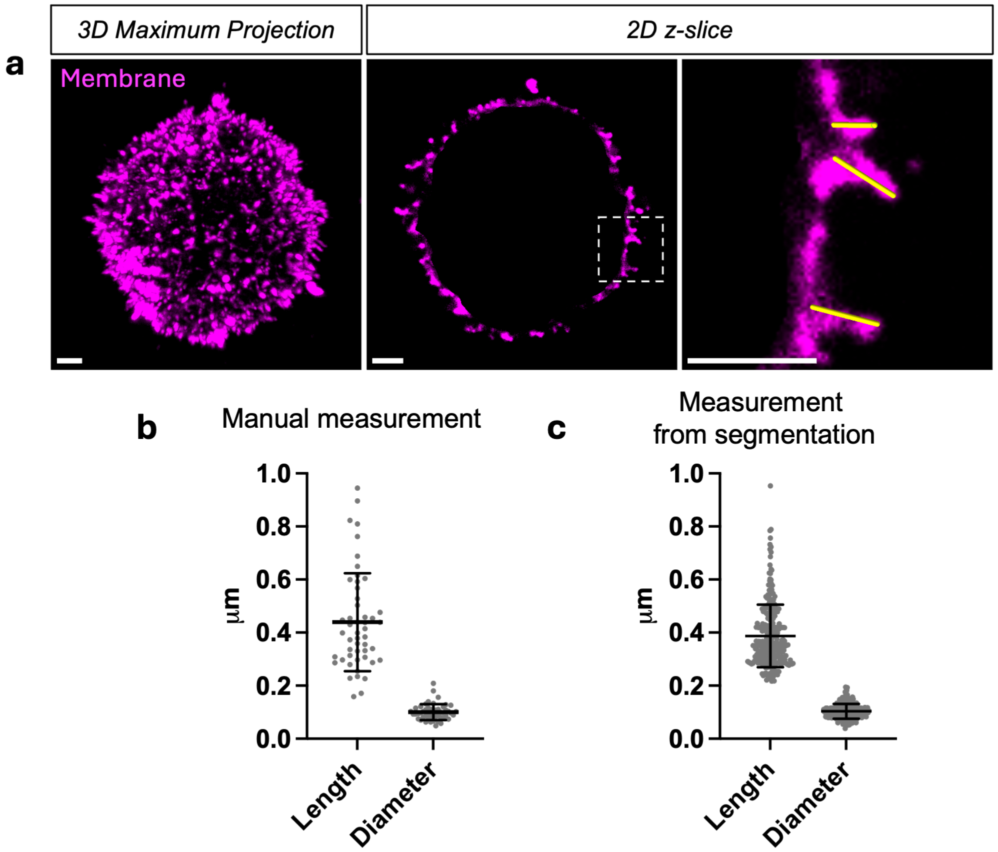


**Figure S3. Manual measurement of microvilli.**

a) 3D maximum projection image and 2D z-slice image of OT-I T cell labeled with WGA-CF633. Magnified image of the region within the white box. The yellow bars show the manual measurements of the microvilli. b) Manual measurement of the microvilli length and diameter of the cell in (a) (*n* = 1 cell, 50 microvilli). c) Measurement of the microvilli length and diameter of the cell in (a) using the segmentation from NanoMAP (*n* = 278 microvilli). Manual measurements were performed with UCSF ChimeraX[74,75]. Scale bar = 1 µm.

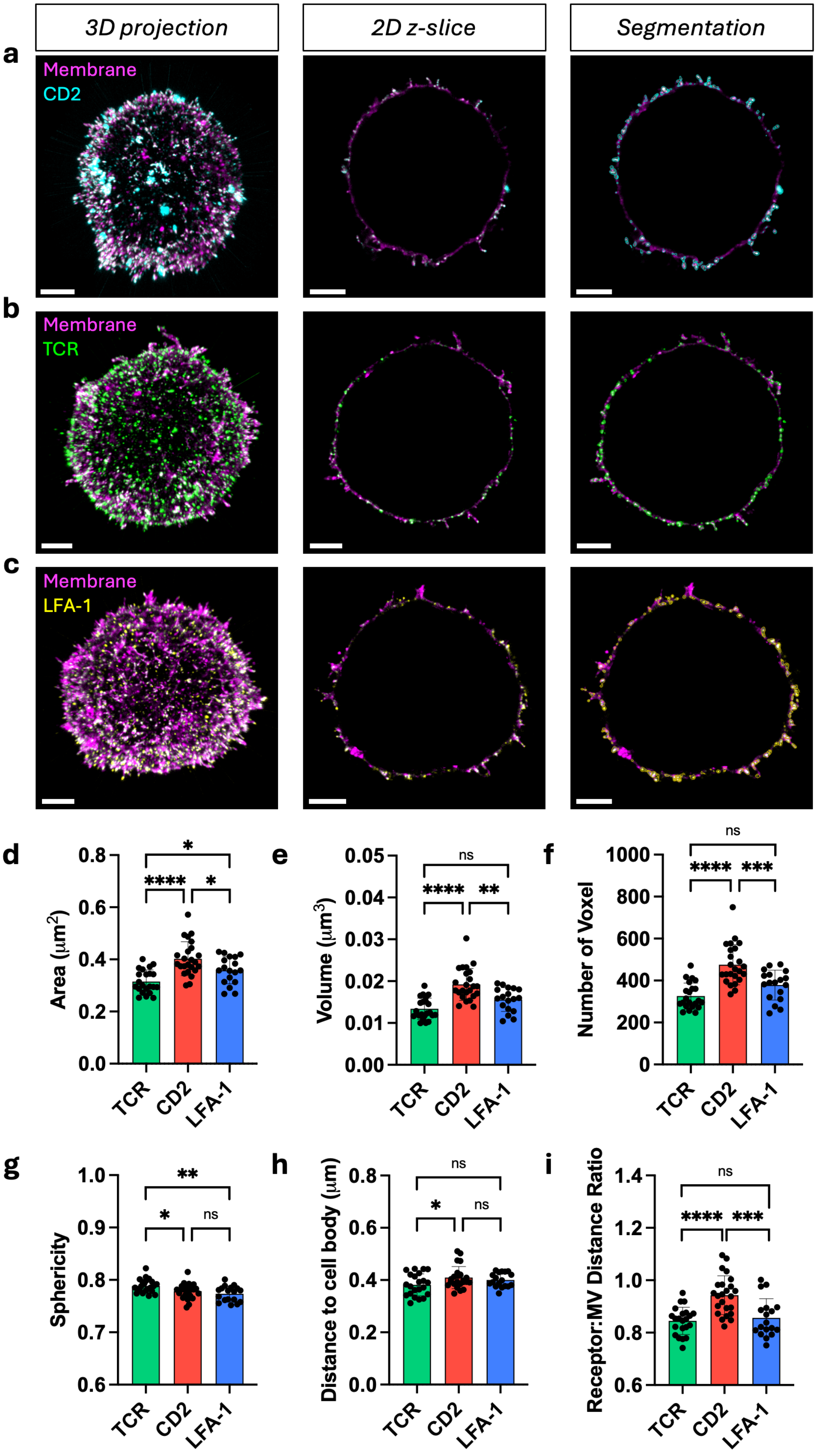
3D projection
2D z-slice
Segmentation
a
Membrane
CD2
b
Membrane
TCR
c
Membrane
LFA-1
d
Area (μm2)
TCR
CD2
LFA-1
e
Volume (μm3)
f
Number of Voxel
g
Sphericity
h
Distance to cell body (μm)
i
Receptor:MV Distance Ratio

**Figure S4. Patch segmentation and characterization**

a-c) Post-expansion images of OT-I T cell in 3D maximum projection, 2D z-slice, and with patch segmentation labeled with WGA-CF633 staining and (a) anti-CD2-AF555, (b) anti-TCR-AF488, and (c) anti-LFA-1-AF555. d) Mean area of TCR, CD2, and LFA-1 patches (CD2 vs. LFA-1: *$p = 0.0328$; CD2 vs. TCR: ****$p < 0.0001$; TCR vs. LFA-1: *$p = 0.0423$). e) Mean volume of TCR, CD2, and LFA-1 patches (CD2 vs. LFA-1: **$p = 0.0015$; CD2 vs. TCR: ****$p < 0.0001$; TCR vs. LFA-1: $p = 0.0754$; ns = not significant). f) Mean number of voxels of TCR, CD2, and LFA-1 patches (CD2 vs. LFA-1: ***$p = 0.0006$; CD2 vs. TCR: ****$p < 0.0001$; TCR vs. LFA-1: $p = 0.0993$; ns = not significant). g) Mean sphericity of TCR, CD2, and LFA-1 patches ranging from 0 to 1, where sphericity=1 indicates a perfect sphere (CD2 vs. LFA-1: $p = 0.4946$; CD2 vs. TCR: *$p = 0.0462$; TCR vs. LFA-1: **$p = 0.0036$; ns = not significant). h) Mean patch distances measured from the cell-body reference surface per cell (CD2 vs. LFA-1: $p = 0.7438$; CD2 vs. TCR: *$p = 0.0481$; TCR vs. LFA-1: $p = 0.2862$; ns = not significant). i) Mean receptor:MV distance ratio calculated as the maximal distance of each patch from the cell-body reference surface divided by the microvillus length (CD2 vs. LFA-1: ***$p = 0.0003$; CD2 vs. TCR: ****$p < 0.0001$; TCR vs. LFA-1: $p = 0.8528$; ns = not significant). In (d-i), data are presented as mean ± S.D. and statistical significance was determined using ordinary one-way ANOVA followed by Tukey's multiple comparisons test. Sample sizes in (d-i): TCR ($n = 22$ cells from N = 6 mice), CD2 ($n = 24$ cells from N = 6 mice), and LFA-1 ($n = 18$ cells from N = 4 mice). Scale bar = 2 μm.

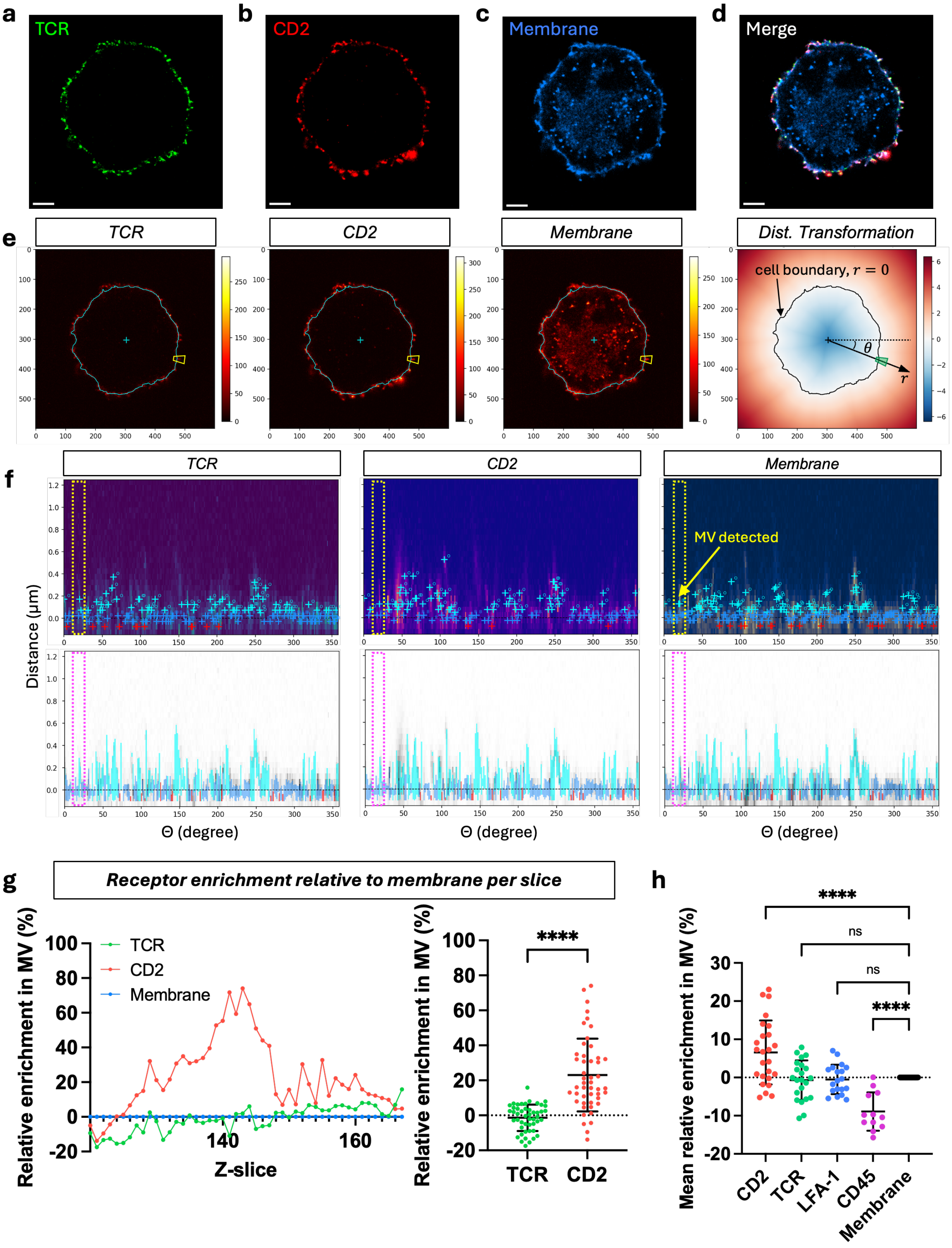

a
TCR
b
CD2
c
Membrane
d
Merge
e
TCR
CD2
Membrane
Dist. Transformation
cell boundary, $r = 0$
$\theta$
$r$
f
TCR
CD2
Membrane
MV detected
Distance (μm)
Θ (degree)
g
Receptor enrichment relative to membrane per slice
Relative enrichment in MV (%)
TCR
CD2
Membrane
Z-slice
****
h
Mean relative enrichment in MV (%)
****
ns
ns
****
CD2
TCR
LFA-1
CD45
Membrane

**Figure S5. Membrane-normalized receptor enrichment analysis in microvilli**

a-d) Post-expansion 2D z-slice images of OT-I T cell labeled with (a) anti-TCR-AF488, (b) anti-CD2-AF555, (c) WGA-CF633 (blue) and (d) all channels merged. Scale bar = 2 µm. e) Images of the same z-slice with the cell boundary (cyan) overlaid on the fluorescence intensity of the TCR, CD2, and membrane channels. The yellow box highlights an example membrane region containing a microvillus. The distance transformation of the z-slice shows the cell boundary (black), cell centroid (black cross), and the polar coordinates $(\theta, r)$ for the highlighted example membrane region. f) Top: Representative peak-detection results from a single 2D z-slice of a cell. Normalized polar intensity maps of the TCR, CD2, and membrane channels are plotted as a function of angular position $\theta$ (x-axis) and distance from the cell boundary $r$ (y-axis). Peaks detected for each angular step are mapped and classified by cell boundary (blue cross), outside boundary (cyan cross) or inside boundary (red cross). The dotted yellow boxes correspond to the membrane region indicated by the yellow box from e). Bottom: Microvilli regions (cyan) and cell-body regions (blue and red), defined from the membrane channel, overlaid with the normalized fluorescence intensity for each individual channel (grayscale). g) Relative outside intensity enrichment of TCR and CD2 in the microvilli calculated for each z-slice from a single cell: (left) enrichment of CD2 and TCR each z-slice and (right) mean ± S.D. across all 2D z-slices. Statistical significance tested using paired two-tailed t-test (****$p$ < 0.0001, $n$ = 48 slices from N = 1 cell). The dashed line represents the membrane baseline at 0 with positive values indicating microvilli localization outside the boundary. h) Average relative outside intensity enrichment of CD2 ($n$ = 25 cells from N = 6 mice), TCR ($n$ = 22 cells from N = 6 mice), LFA-1 ($n$ = 19 cells from N = 4 mice), and CD45 ($n$ = 12 cells from N = 2 mice) relative to the normalized membrane channel ($n$ = 33 cells from N = 6 mice). Data are presented as mean ± S.D. Statistical significance was determined using ordinary one-way ANOVA followed by Dunnett's comparisons test against membrane as control (membrane vs. CD2: ****$p$ < 0.0001; membrane vs. TCR: $p$ = 0.9715; membrane vs. LFA-1: $p$ = 0.9932; membrane vs. CD45: ****$p$ < 0.0001; ns = not significant).

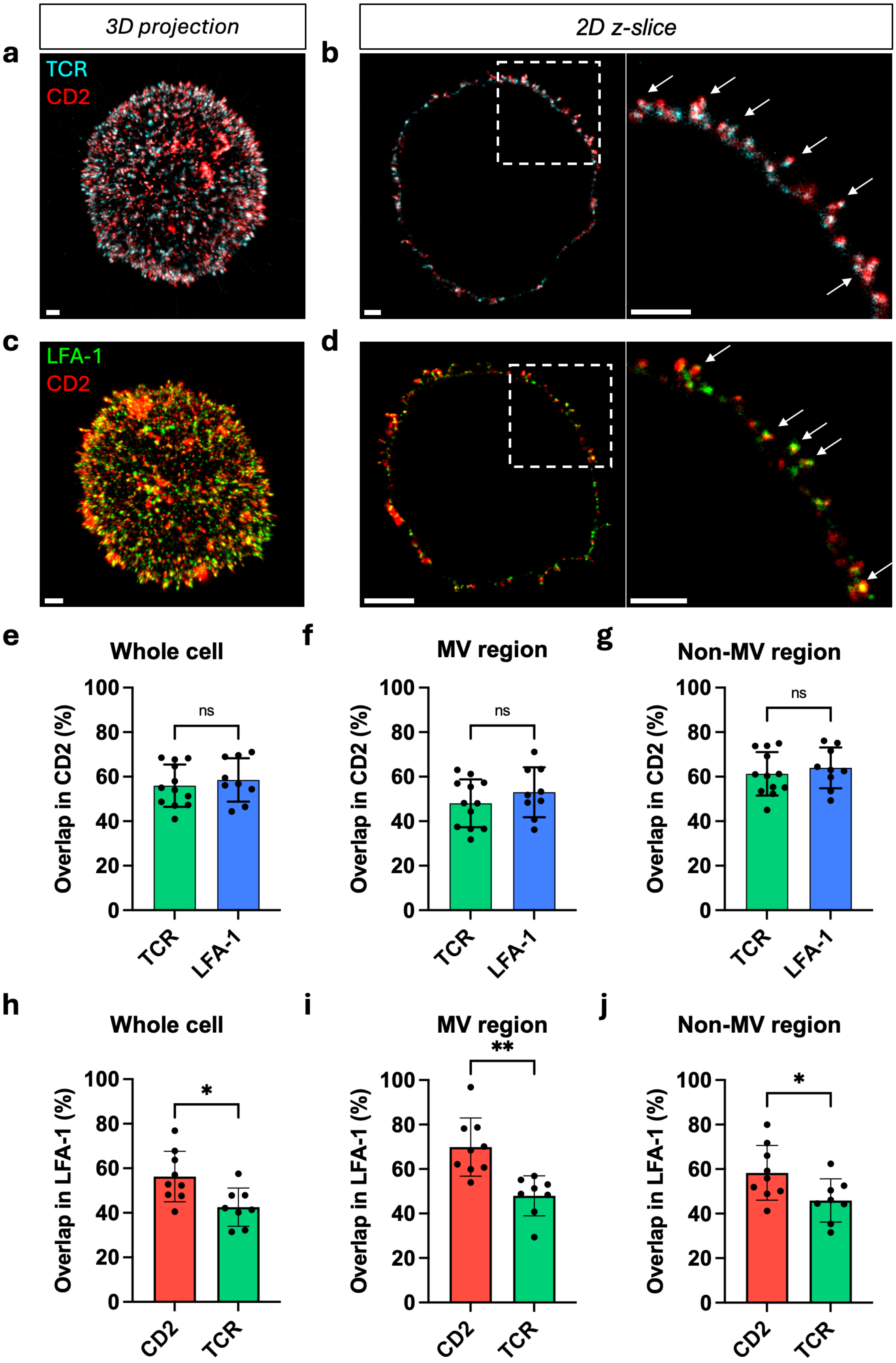
3D projection
2D z-slice
a
TCR
CD2
b
c
LFA-1
CD2
d
e
Whole cell
f
MV region
g
Non-MV region
Overlap in CD2 (%)
100
80
60
40
20
0
ns
TCR
LFA-1
h
Whole cell
i
MV region
j
Non-MV region
Overlap in LFA-1 (%)
*
**
CD2
TCR

**Figure S6. CD2 shows similar degree of association with TCR and LFA-1 patches.**

a-b) Post-expansion image of OT-I T cell labeled with anti-TCR-AF488 (cyan) and anti-CD2-AF555 (red) in (a) 3D maximum projection and (b) 2D z-slice with a magnified image of the region within the white box. White arrows are pointing at the microvilli tips. c-d) Post-expansion image of a T cell labeled with anti-CD2-AF555 (red) and anti-LFA-1-AF488 (green) in (c) 3D maximum projection and (d) 2D z-slice with a magnified image of the region within the white box. White arrows are pointing at the microvilli tips. e-g) Colocalization of TCR and LFA-1 patches with CD2 patches in the (e) whole T cell membrane, (f) MV region, and (g) non-MV region (whole cell: $p = 0.5483$; MV region: $p = 0.3231$; non-MV region: $p = 0.5311$; ns = not significant). Sample sizes: TCR ($n = 12$ cells from N = 3 mice), and LFA-1 ($n = 9$ cells from N = 2 mice). h-j) Colocalization of TCR and CD2 patches with LFA-1 patches in the (h) whole T cell membrane, (i) MV region, and (j) non-MV region (whole cell: $*p = 0.0127$; MV region: $**p = 0.0011$; non-MV region: $*p = 0.0350$). Sample sizes: CD2 ($n = 9$ cells from N = 2 mice), and TCR ($n = 8$ cells from N = 2 mice). In (e-j), data are presented as mean ± S.D. Statistical significance tested using unpaired two-tailed Welch's t-test. Scale bar = 1 μm.

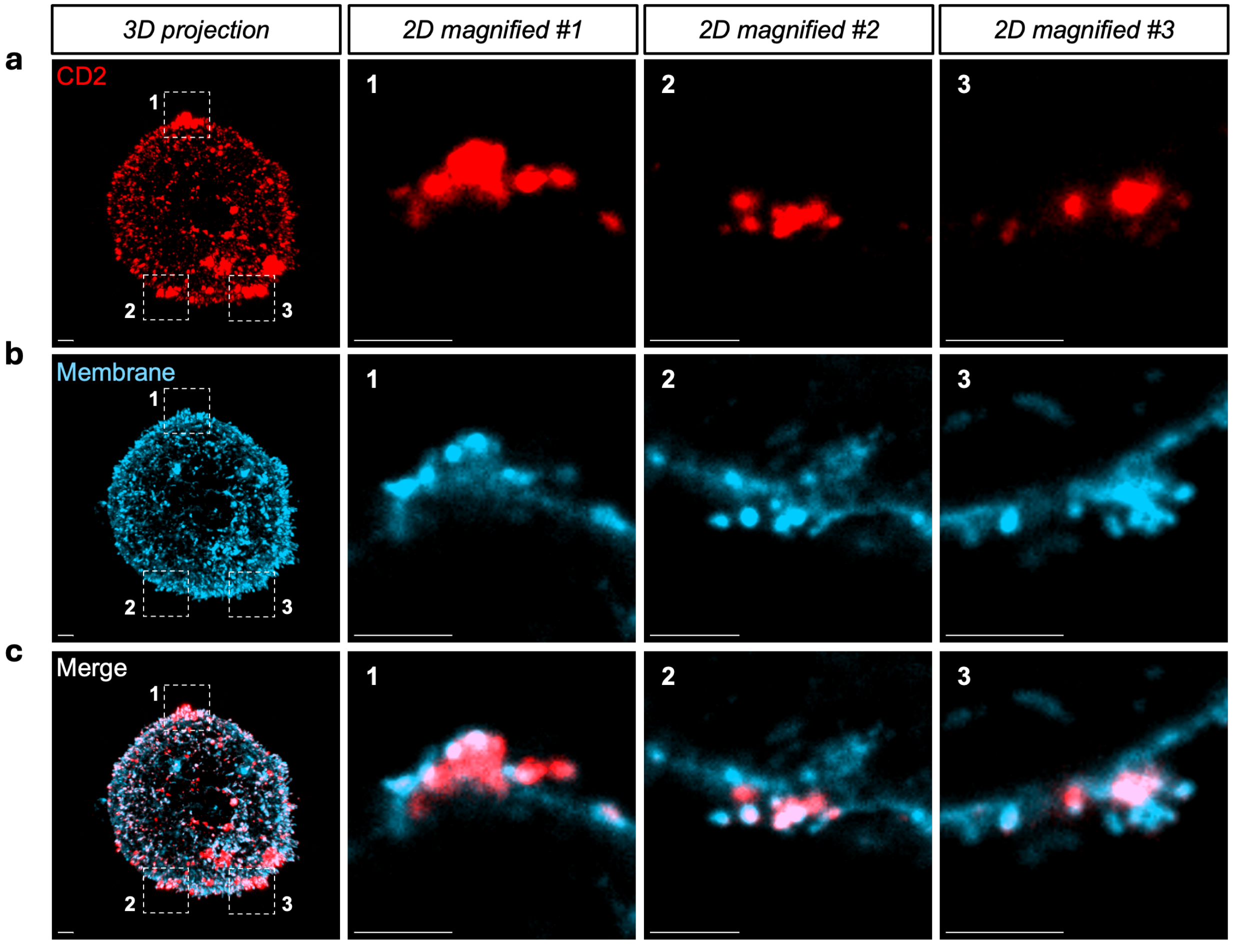


**Figure S7. Large CD2 patch localized on T cell membrane.**

a) Post-expansion 3D image of OT-I T cell labeled with anti-CD2 (red) and magnified 2D z-slice of the region within the white boxes labeled #1, #2, and #3. b) Post-expansion 3D image of the T cell in (a) stained with WGA-CF633 (blue). Regions within the white boxes #1, #2, and #3 are magnified in 2D z-slice images. c) Merged images from (a) and (b). Scale bar = 1 μm.

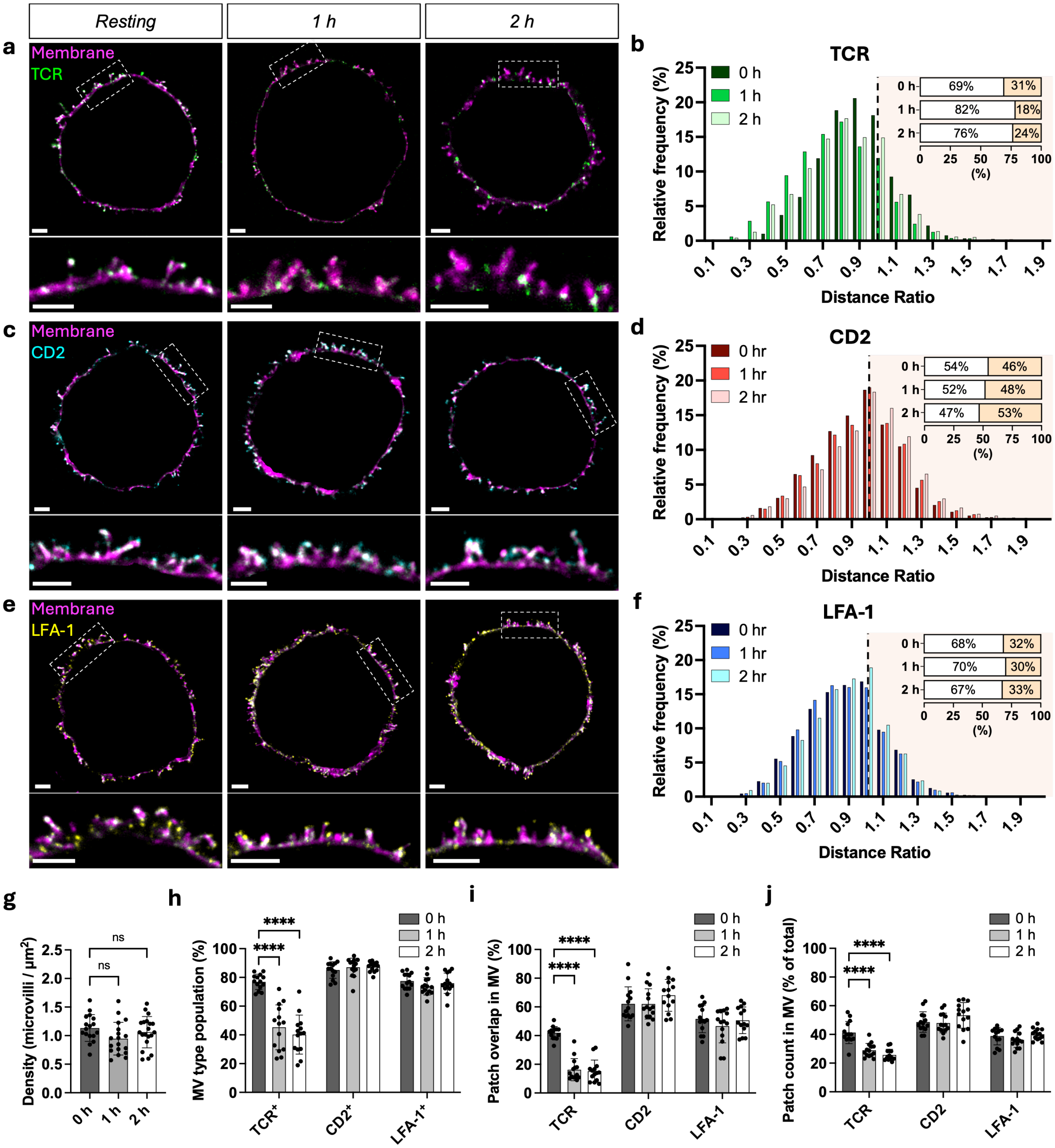

Resting
1 h
2 h
a
Membrane
TCR
b
TCR
0 h
1 h
2 h
0 h 69% 31%
1 h 82% 18%
2 h 76% 24%
(%)
Relative frequency (%)
Distance Ratio
c
Membrane
CD2
d
CD2
0 hr
1 hr
2 hr
0 h 54% 46%
1 h 52% 48%
2 h 47% 53%
(%)
Relative frequency (%)
Distance Ratio
e
Membrane
LFA-1
f
LFA-1
0 hr
1 hr
2 hr
0 h 68% 32%
1 h 70% 30%
2 h 67% 33%
(%)
Relative frequency (%)
Distance Ratio
g
Density (microvilli / µm²)
ns
ns
0 h
1 h
2 h
h
MV type population (%)
****
****
TCR⁺
CD2⁺
LFA-1⁺
i
Patch overlap in MV (%)
****
****
TCR
CD2
LFA-1
j
Patch count in MV (% of total)
****
****
TCR
CD2
LFA-1

**Figure S8. Redistribution of TCR and adhesion proteins of resting and activated T cells**

a) Post-expansion 2D z-slice image (top) and magnified image of the region within the white box (bottom) of OT-I T cell labeled with anti-TCR-AF488 (green) and WGA-CF633 (magenta) at resting (0 h), 1 h, and 2 h activation after co-incubation with pMHC-coated beads. b) Relative frequency distribution of TCR patch distance ratios normalized to microvillus length. c) Post-expansion 2D z-slice image (top) and magnified image (bottom) of a T cell labeled with anti-CD2-AF555 (cyan) and WGA-CF633 (magenta) at resting (0 h), 1 h, and 2 h activation after co-incubation with pMHC-coated beads. d) Relative frequency distribution of CD2 patch distance ratios normalized to microvillus length. e) Post-expansion 2D z-slice image (top) and magnified image (bottom) of a T cell labeled with anti-LFA-1-AF555 (yellow) and WGA-CF633 (magenta) at resting (0 h), 1 h, and 2 h activation after co-incubation with pMHC-coated beads. f) Relative frequency distribution of LFA-1 patch distance ratios normalized to microvillus length. In (b), (d), (f), Inset: distribution (%) with distance ratio <1 (below the tip) and ≥1 (at or beyond the tip). g) Mean density of microvilli per area (0 h vs. 1 h: $p = 0.0799$; 0 h vs. 2 h: $p = 0.6242$; ns = not significant). Sample sizes in (g): 0 h ($n = 17$ cells from N = 4 mice), 1 h ($n = 18$ cells from N = 4 mice), and 2 h ($n = 20$ cells from N = 4 mice). h) Population of microvilli containing TCR, CD2, and LFA-1 patches per cell at resting (0 h), 1 h, and 2 h activation (TCR 0 h vs. 1 h: ****$p < 0.0001$; TCR 0 h vs. 2 h: ****$p < 0.0001$; CD2 0 h vs. 1 h: $p = 0.9998$; CD2 0 h vs. 2 h: $p = 0.2792$; LFA-1 0 h vs. 1 h: $p = 0.3417$; LFA-1 0 h vs. 2 h: $p = 0.9557$; ns = not significant). i) Mean occupancy (voxel overlap) of TCR, CD2, and LFA-1 in the MV region per cell at resting (0 h), 1 h, and 2 h activation (TCR 0 h vs. 1 h: ****$p < 0.0001$; TCR 0 h vs. 2 h: ****$p < 0.0001$; CD2 0 h vs. 1 h: $p = 0.5433$; CD2 0 h vs. 2 h: $p = 0.4237$; LFA-1 0 h vs. 1 h: $p = 0.2218$; LFA-1 0 h vs. 2 h: $p = 0.7538$; ns = not significant). j) Fraction of the number of patches in the MV region to the total number of patches in the whole cell at resting (0 h), 1 h, and 2 h activation. (TCR 0 h vs. 1 h: ****$p < 0.0001$; TCR 0 h vs. 2 h: ****$p < 0.0001$; CD2 0 h vs. 1 h: $p = 0.9623$; CD2 0 h vs. 2 h: $p = 0.1985$; LFA-1 0 h vs. 1 h: $p = 0.3254$; LFA-1 0 h vs. 2 h: $p = 0.7372$; ns = not significant). In (g-j), statistical significance was determined using ordinary one-way ANOVA with Dunnett's comparisons test against 0 h. Sample sizes in (a-f) and (h-j): $n = 14$ cells from N = 3 mice for all conditions. Scale bar = 1 μm.

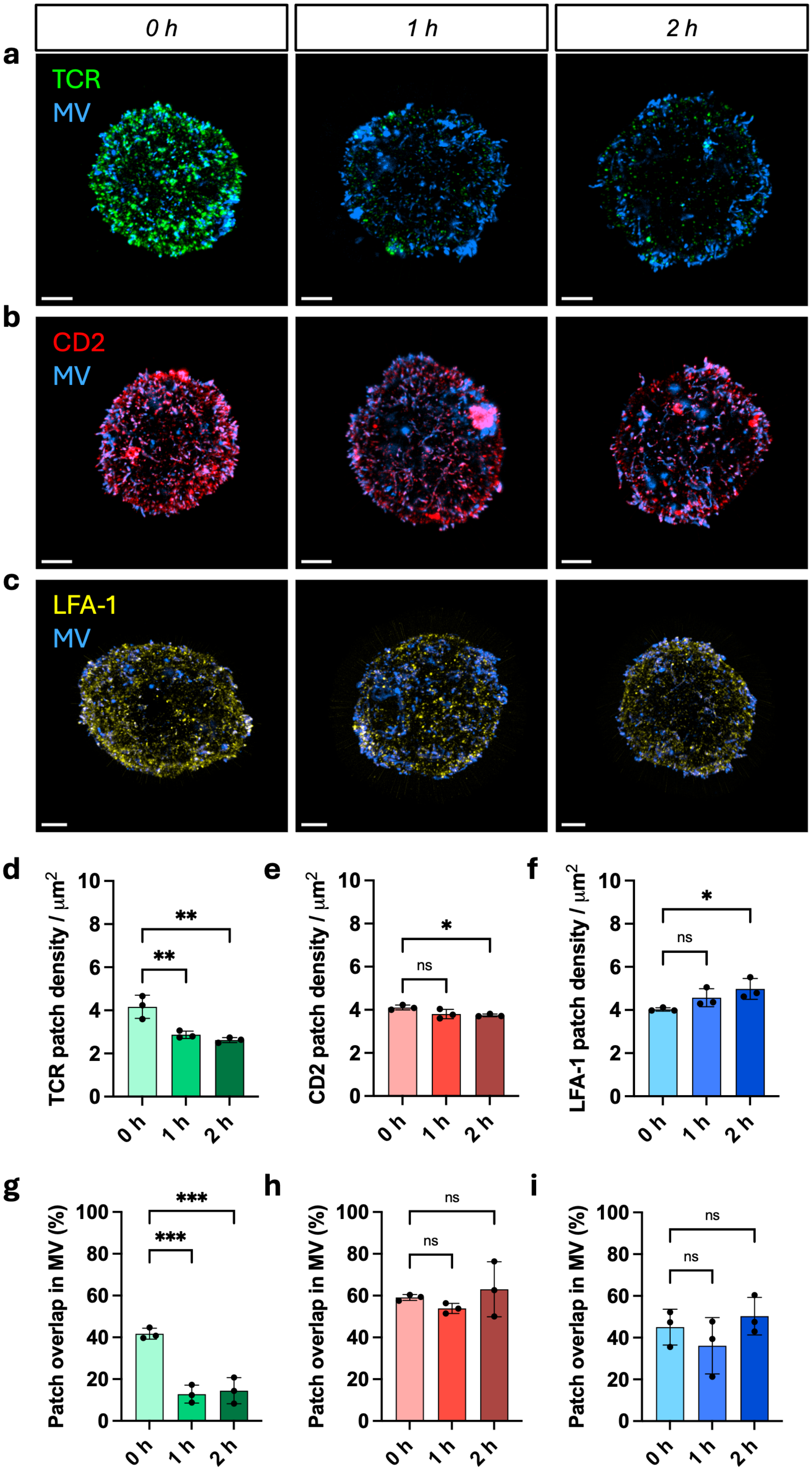

0 h
1 h
2 h
a
TCR
MV
b
CD2
MV
c
LFA-1
MV
d
TCR patch density / μm2
**
**
e
CD2 patch density / μm2
ns
*
f
LFA-1 patch density / μm2
ns
*
g
Patch overlap in MV (%)
***
***
h
Patch overlap in MV (%)
ns
ns
i
Patch overlap in MV (%)
ns
ns

**Figure S9. TCR depletion from microvilli after T cell:MC38 co-incubation.**

a-c) Post-expansion image of isolated T cell after co-incubation with MC38 tumor cell for 0 h, 1 h, and 2 h in 3D maximum projection. T cell is labeled with a) anti-TCR (green), b) anti-CD2 (red), c) anti-CD11a (yellow) and a-c) WGA (blue) for membrane staining. d-f) Mean patch density of d) TCR, e) CD2, and f) LFA-1 per cell after 0 h, 1 h, and 2 h co-incubation with MC38 tumor cell. (TCR 0 h vs. 1 h: $^{**}p = 0.0055$; TCR 0 h vs. 2 h: $^{**}p = 0.0023$; CD2 0 h vs. 1 h: $p = 0.0774$; CD2 0 h vs. 2 h: $^{*}p = 0.0399$; LFA-1 0 h vs. 1 h: $p = 0.2010$; LFA-1 0 h vs. 2 h: $^{*}p = 0.0351$; ns = not significant). g-i) Mean occupancy (voxel overlap) of g) TCR, h) CD2, and i) LFA-1 in the MV region per cell after 0 h, 1 h, and 2 h co-incubation with MC38 tumor cell. (TCR 0 h vs. 1 h: $^{***}p = 0.0005$; TCR 0 h vs. 2 h: $^{***}p = 0.0007$; CD2 0 h vs. 1 h: $p = 0.6422$; CD2 0 h vs. 2 h: $p = 0.7743$; LFA-1 0 h vs. 1 h: $p = 0.5211$; LFA-1 0 h vs. 2 h: $p = 0.7757$; ns = not significant). Sample sizes: $n = 3$ cells from N = 1 mice. Data are presented as mean ± S.D. Statistical significance tested using ordinary one-way ANOVA with Dunnett's comparisons test compared against 0 h. Scale bar = 1 μm.

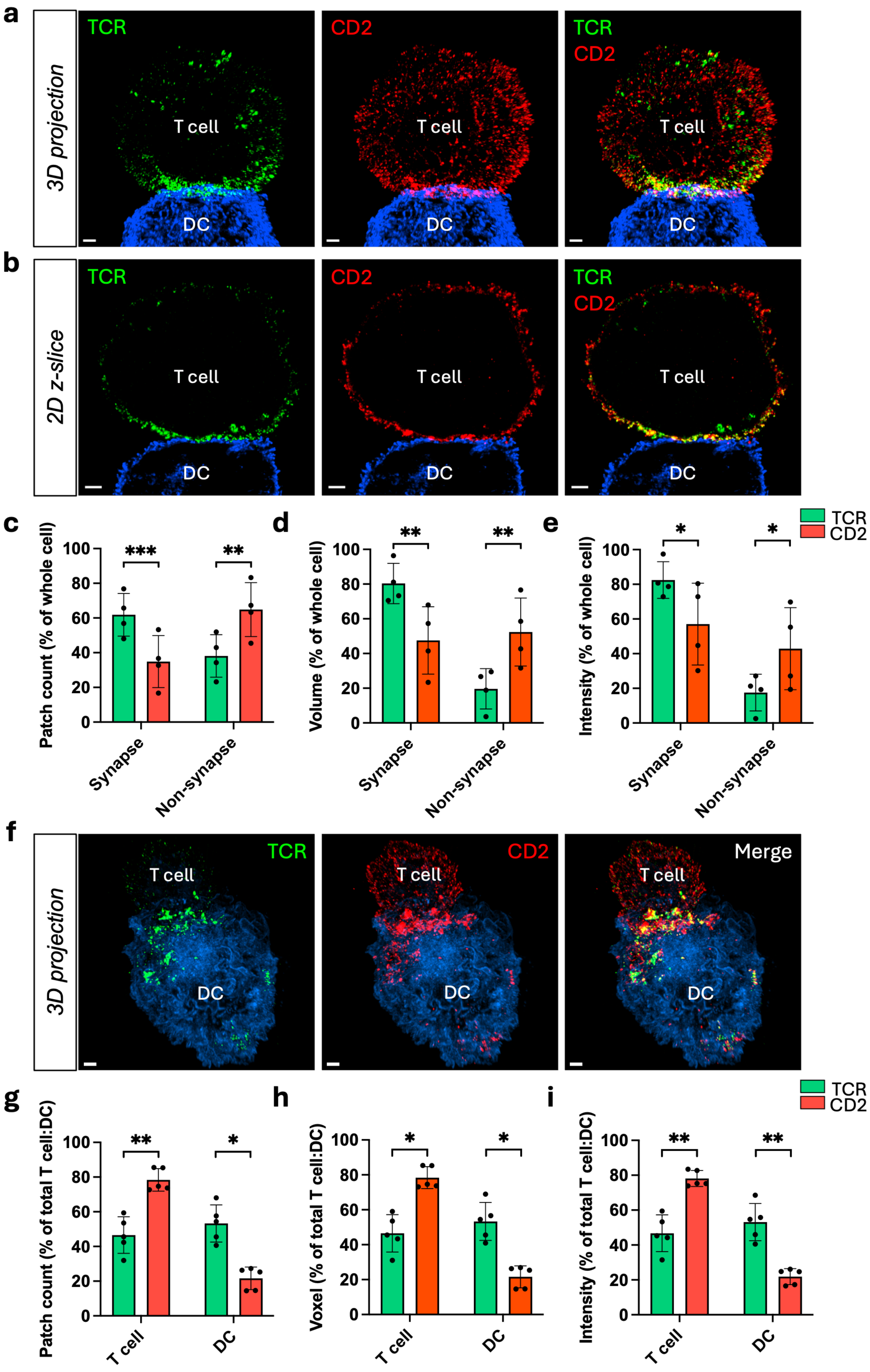

a
3D projection
TCR
T cell
DC
CD2
T cell
DC
TCR
CD2
T cell
DC
b
2D z-slice
TCR
T cell
DC
CD2
T cell
DC
TCR
CD2
T cell
DC
c
Patch count (% of whole cell)
***
**
Synapse
Non-synapse
d
Volume (% of whole cell)
**
**
Synapse
Non-synapse
e
Intensity (% of whole cell)
*
*
Synapse
Non-synapse
TCR
CD2
f
3D projection
TCR
T cell
DC
CD2
T cell
DC
Merge
T cell
DC
g
Patch count (% of total T cell:DC)
**
*
T cell
DC
h
Voxel (% of total T cell:DC)
*
*
T cell
DC
i
Intensity (% of total T cell:DC)
**
**
T cell
DC
TCR
CD2

**Figure S10. TCR and CD2 exhibit different migration patterns and patch shedding on DC during synapse formation.**

a-b) Post-expansion image of OT-I T cell conjugated with a DC for 10 m in (a) 3D maximum projection and (b) 2D z-slice. T cell is labeled with anti-TCR-AF488 (green), anti-CD2-AF555 (red) and WGA-CF633 (blue) for DC membrane staining. c) Fraction of the patch count in the synapse or non-synapse region to the total patch count in the whole cell (TCR vs. CD2 synapse: ***$p$ = 0.0007; non-synapse: **$p$ = 0.0010). d) Fraction of the patch volume in the synapse or non-synapse region to the total patch volume in the whole cell (TCR vs. CD2 synapse: **$p$ = 0.0074; non-synapse: **$p$ = 0.0076). e) Fraction of the intensity sum in the synapse or non-synapse region to the total intensity sum in the whole cell (TCR vs. CD2 synapse: *$p$ = 0.0488; non-synapse: *$p$ = 0.0494). Sample sizes: $n$ = 4 cells from N = 2 mice. f) Post-expansion image of T cell conjugated with a DC for 10 m in 3D maximum projection. T cell is labeled with anti-TCR-AF488 (green), anti-CD2-AF555 (red) and WGA-CF633 (blue) for DC membrane staining. g) Fraction of the patch count in the T cell or DC membrane to the total patch count in the whole T cell:DC conjugate (TCR vs. CD2 T cell: **$p$ = 0.0098; DC: *$p$ = 0.0104). h) Fraction of the voxel number in the T cell or DC membrane to the total voxel number in the whole T cell:DC conjugate (TCR vs. CD2 T cell: *$p$ = 0.0102; DC: *$p$ = 0.0108). i) Fraction of the intensity sum in the T cell or DC membrane to the total intensity sum in the whole T cell:DC conjugate (TCR vs. CD2 T cell: **$p$ = 0.0077; DC: **$p$ = 0.0081). Sample sizes: $n$ = 5 cells from N = 2 mice. Data are presented as mean ± S.D. Statistical significance tested using two-tailed paired t-test comparing TCR and CD2 measurements within the same cells. Scale bar = 1 μm.

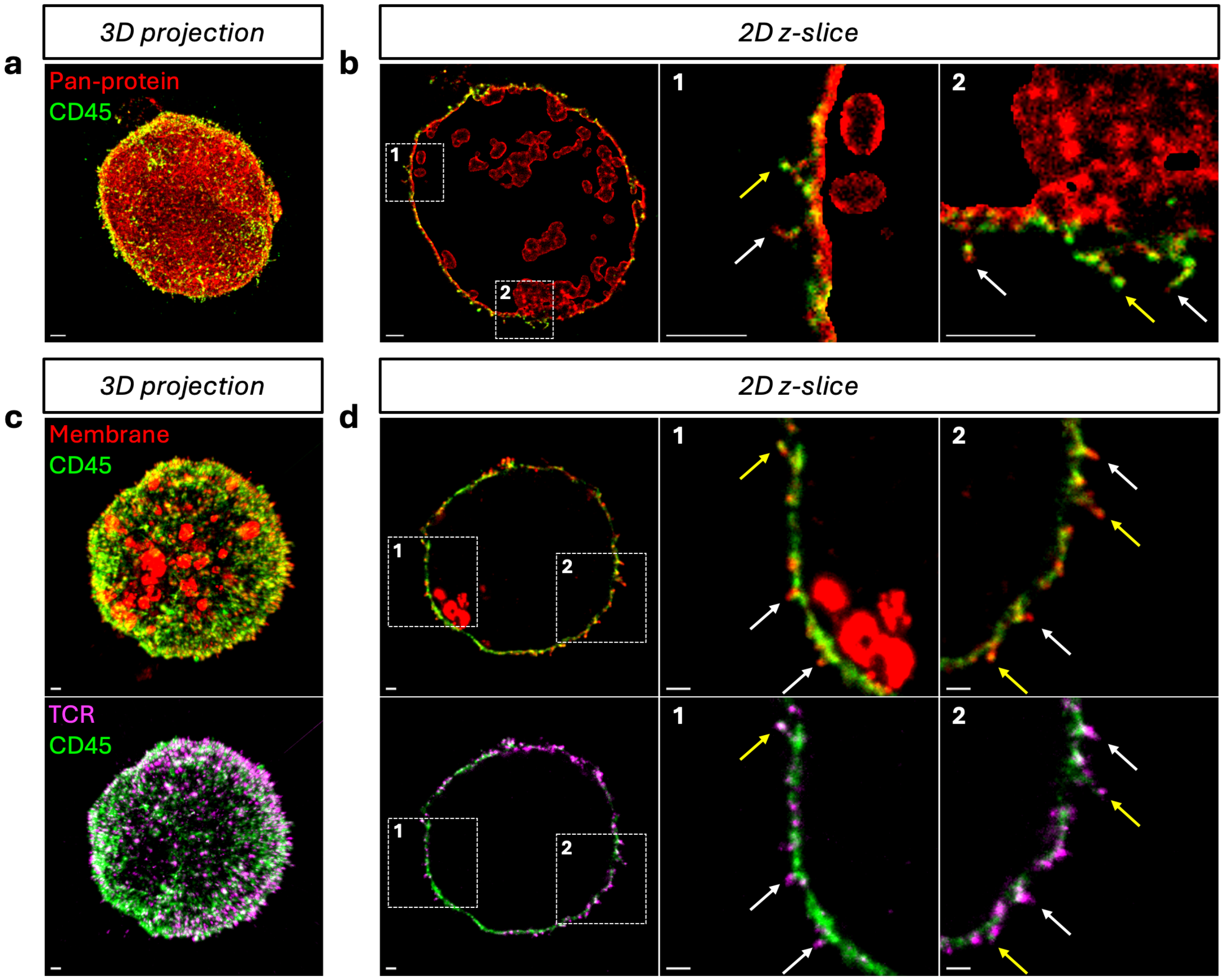


**Figure S11. CD45 patches exhibit partial exclusion on the microvilli**

a) 3D image of OT-I T cell fully expanded in $H_2O$ and labeled with anti-CD45 (green) and NHS-ester Cy5 for pan-protein staining (red). b) 2D z-slice image of (a) and magnified image of the region within the white boxes. c) 3D images of T cell expanded in 1x PBS with (top) anti-CD45 and WGA-CF633 and (bottom) anti-CD45 and anti-TCR labeling. d) 2D z-slice images of (c) and magnified images of the region within the white boxes. In (b) and (d), white arrows are pointing at CD45 exclusion on the microvilli and yellow arrows are pointing at CD45 partial exclusion or localization on the microvilli. Scale bar = 0.5 µm.

**Movie S1. 3D visualization of the synaptic cleft between a T cell-MC38 tumor cell conjugate at ~35 nm resolution**

3D rendering of a fixed OT-I T cell-MC38 tumor cell conjugate, fully expanded in $H_2O$ to achieve ~35 nm resolution. The T cell-MC38 tumor cell conjugate is labeled with anti-CD45 (green) and NHS-ester (magenta) for pan-protein staining. Rendered T cell and MC38 cell surfaces illustrate the architecture of the synaptic cleft in an *en face* view. The rendering displays the individual 2D z-slices that make up the 3D image. The movie shows a single fixed and expanded T cell-MC38 cell conjugate and contains no temporal information with apparent motion reflecting changes in rendered viewpoint only.

**Movie S2. TCR organization and membrane topology on the surface of naïve and effector T cells**

Rotating 3D rendering of fixed and expanded naïve and effector OT-I T cells demonstrating the distribution of TCR and microvillar structure of the cell membrane. T cells were labeled with anti-TCR and WGA for membrane staining. Each rendering shows a single fixed, expanded cell, with rotation reflecting changes in viewing angle only.

**Movie S3. Visualization and segmentation of microvilli and TCR-CD2 patches in a fixed and expanded OT-I T Cell**

3D rendering of a fixed and expanded OT-I T cell labeled with anti-CD2 (red), anti-TCR (green), and WGA (blue) for membrane staining. The rendering shows a 2D z-slice of the cell, demonstrating how the MV region is isolated from membrane signal by removing the cell body (non-MV region) signal, which enables segmentation of microvilli surfaces. The magnified views show segmented CD2 and TCR patches overlaid with microvilli surfaces. The movie shows a single fixed and expanded sample. No temporal information is given, and motions reflect changes in rendering viewpoint only.

**Supplementary Table 1.** Primary antibodies for pre- and post-ExM immunostaining

| Host | Target | Vendor | Clone | Catalog # |
|---|---|---|---|---|
| Rat | CD11a | Biolegend | H155-78 | 141002 |
| Rat | CD2 | Biolegend | RM2-5 | 100102 |
| Rabbit | CD2 | Abcam | EPR21825 | ab219411 |
| Rabbit | CD28 | Abcam | EPR22076 | ab243228 |
| Rat | CD4 | Biolegend | RM4-5 | 100506 |
| Rat | CD45 | Biolegend | 30-F11 | 103102 |
| Chicken | CD62L | R&D Systems | - | AF576 |
| Rat | CD8α | Biolegend | 53-6.7 | 100702 |
| Rat | PD-1 | Biolegend | RMP1-30 | 109101 |
| Armenian hamster | TCRβ | Biolegend | H57-597 | 109215 |
| Armenian hamster | TCRβ | Biolegend | H57-597 | 109238 |
| Mouse | Lck | Biolegend | A17013D | 601001 |
| Mouse | ZAP70 | Biolegend | 1E7.2 | 313402 |
| Rabbit | β-actin | Proteintech | - | 20536-1-AP |
| Rabbit | CD2 | Abcam | EPR27426-17 | ab314761 |
| Rabbit | CD3ε | Abcam | - | ab5690 |
| Rabbit | CD45 | Abcam | - | ab10558 |

**Supplementary Table 2.** Secondary antibodies for pre- and post-ExM immunostaining

| Host | Target | Conjugate | Vendor | Catalog # |
|---|---|---|---|---|
| Goat | Rabbit | DyLight 488 | Invitrogen | 35552 |
| Goat | Rabbit | DyLight 550 | Invitrogen | 84541 |
| Goat | Arm. hamster | Alexa Fluor 488 | Invitrogen | A78963 |
| Donkey | Rat | Alexa Fluor 488 | Invitrogen | A21208 |
| Donkey | Rat | Alexa Fluor 555 | Invitrogen | A48270 |
| Donkey | Rat | Alexa Fluor 647 | Invitrogen | A48272 |
| Donkey | Mouse | Alexa Fluor 488 | Invitrogen | A32766 |
| Donkey | Mouse | Alexa Fluor 555 | Invitrogen | A31570 |
| Donkey | Goat | Alexa Fluor 488 | Invitrogen | A32814 |
| Donkey | Goat | Alexa Fluor 555 | Invitrogen | A32816 |
| Donkey | Chicken | Alexa Fluor 555 | Invitrogen | A78949 |

**Supplementary Table 3.** Fluorescent labels for pre- and post-ExM staining

| Staining Agent | Vendor | Catalog # |
|---|---|---|
| Wheat Germ Agglutinin (WGA) CF®633 | Biotium | 29024 |
| NHS ester-Cy5 | Cytiva | PA15101 |
| DAPI | Thermo Scientific | 62248 |

**Supplementary Table 4.** List of reagents for ExM

| Reagent | Vendor | Catalog # |
|---|---|---|
| N,N-dimethylacrylamide | Sigma Aldrich | 274135 |
| Sodium acrylate | AstaTech | H10710 |
| Acrylamide | Sigma Aldrich | A8887 |
| N,N′Methylenebisacrylamide | Sigma Aldrich | M7279 |
| Sodium chloride | Sigma Aldrich | S6191 |
| N,N,N′,N′Tetramethylethylenediamine | Sigma Aldrich | T9281 |
| Methacrolein | Sigma Aldrich | 133035 |
| 2-Hydroxy-4′-(2-hydroxyethoxy)-2-methylpropiophenone | Sigma Aldrich | 106797-53-9 |
| Urea | Sigma Aldrich | U5378 |
| Glycine | Sigma Aldrich | G8898 |
| 0.5M EDTA | VWR | BDH78301 |
| Tris base | Fischer Scientific | BP152-1 |
| Phosphate Buffered Saline, 10x Solution | Sigma-Aldrich | 11666789001 |
| Ultrapure SSC, 20X | Thermo Scientific | 15557044 |
| Tween 20™, Ultrapure, Thermo Scientific™ | Thermo Scientific | J20605.AP |
| Triton™ X-100, BioXtra | Sigma-Aldrich | T9284-100ML |
| 365nm 300W LED UV Curing Lamp | Somesino (via Amazon) | sin-002 |

**Supplementary Table 5.** Cell culture reagents

| Reagent | Vendor | Catalog # |
|---|---|---|
| RPMI | Thermo Scientific | 11875093 |
| DMEM | Thermo Scientific | 11965092 |
| Fetal bovine serum | Thermo Scientific | A5256701 |
| Penicillin-Streptomycin-Glutamine | Thermo Scientific | 10378016 |
| HEPES-buffered saline | Thermo Scientific | J67502.AE |
| 2-Mercaptoethanol | Thermo Scientific | 21985023 |
| DPBS | Thermo Scientific | 14190250 |
| Bovine Serum Albumin | Sigma-Aldrich | A4503-100G |
| Interleukin-2, human, recombinant (E. coli) | Sigma-Aldrich | 11011456001 |
| Recombinant Murine IL-4 | PeproTech | 214-14 |
| Recombinant Murine GM-CSF | PeproTech | 315-03 |
| EasySep Mouse CD8+ T Cell Isolation Kit | STEMCELL | 19853 |
| EasySep Mouse CD4+ T Cell Isolation Kit | STEMCELL | 19852 |
| EasySep Human CD8+ T Cell Isolation Kit | STEMCELL | 17953 |
| Dynabeads Mouse T-Activator CD3/CD28 | Thermo Fisher | 11453D |
| Dynabeads M-280 Streptavidin | Thermo Fisher | 11205D |
| Human Peripheral Blood Mononuclear Cells | STEMCELL | 70025 |
| ImmunoCult Human CD3/CD28 T Cell Activator | STEMCELL | 10971 |
| ImmunoCult-XF T cell Expansion Medium | STEMCELL | 10981 |

**Supplementary Table 6.** SLB reagents

| Reagent | Vendor | Catalog # |
|---|---|---|
| 1-palmitoyl-2-oleoyl-sn-glycero-3-phosphocholine | Avanti Polar Lipids | 850457 |
| 1,2-dioleoyl-sn-glycero-3-[(N-(5-amino-1-carboxypentyl) iminodiacetic acid) succinyl] (nickel salt) | Avanti Polar Lipids | 790404 |
| 1,2-dioleoyl-sn-glycero-3-phosphoethanolamine-N-(cap biotinyl) (sodium salt) | Avanti Polar Lipids | 870273 |
| 1,2-dioleoyl-sn-glycero-3-phosphoethanolamine-N-[methoxy (polyethylene glycol)-5000] (ammonium salt) | Avanti Polar Lipids | 880230 |
| LiposoFast extruder | Avestin | LF-1 |
| Hellmanex | Sigma-Aldrich (MilliporeSigma) | Z805939 |
| ICAM-1 | Sino Biological | 50440-M08H |
| H-2K$^{b}$–OVA$^{257\text{-}264}$ (SIINFEKL) | NIH Tetramer Core Facility | reagent ID 4143 |
| Nunc™ Lab-Tek™ II Chambered Coverglass | Thermo Scientific | 155382PK |
